# The Intermediate Hippocampus Integrates Shock-Observation and Spatial Information during Observational Fear Memory in Male Rats

**DOI:** 10.64898/2026.01.03.697465

**Authors:** Frédéric Michon Linde, Valeria Gazzola, Christian Keysers

## Abstract

Learning where danger lurks by observing others requires linking another individual’s distress to one’s own representation of space. How the hippocampus implements this mapping is unknown. We recorded Neuropixels activity across the dorsal, intermediate and ventral hippocampus of male rats during contextual fear learning by observation, subsequent rest and recall. Observers showed increased immobility in the context where they had witnessed a conspecific receive footshocks, with substantial inter-individual variability. Pyramidal neurons were recruited by shock observation, with enriched shock-excited cells in the intermediate hippocampus. In the dorsal and intermediate hippocampus, shock-observation responses scaled with baseline place-field firing, revealing a gain-like conjunctive code between observed distress and observer’s position. Following learning, shock-context spatial representations were selectively stabilized in the intermediate hippocampus, and post-learning sharp-wave ripples preferentially reactivated shock-context ensembles, especially in animals expressing contextual freezing.These findings identify candidate hippocampal mechanisms by which socially acquired threat information are mapped onto the observer’s spatial representation and consolidated into memory.

## Introduction

To safely navigate their environment, animals must learn where danger lies. Learning from others’ misfortunes is safer than relying on direct experience. Mounting evidence suggests that rodents orient towards conspecifics in altered affective states^1^ and can exhibit distress when witnessing the distress of others^2–4^. Notably, they show elevated freezing when re-exposed to the context in which they previously witnessed a conspecific receive shocks^3,5^, indicating that they can associate others’ distress with the context in which it was observed.

At the mechanistic level, neurons involved in first-hand distress are activated when rodents witness the distress of others in cingulate Area 24^6^, and interfering with activity in Area 24, the basolateral amygdala, midline thalamic nuclei, or dorsal hippocampus disrupts observational fear learning^3,5–8^. However, these findings leave unresolved a key question: if animals can learn where danger lies by observing others, how is this socially acquired information incorporated into the hippocampal cognitive maps that guide behavior?

The hippocampus is a key structure for forming these cognitive maps that support flexible navigation^9^. During active experience, pyramidal neurons fire in a location-specific manner, collectively forming a spatial representation of the environment that can integrate internal states and task variables^10–17^. Individual experiences are associated with unique patterns of activity across hippocampal populations^18,19^. The stabilization of these representations is linked to learning^20,21^, and their reinstatement contributes to memory retrieval^22–24^. During post-experience rest or sleep, experience-related activity is replayed in a time-compressed manner during sharp-wave ripple (SWR) events, which are causally implicated in memory consolidation^25–31^.

The hippocampus is organized along its septotemporal axis into subregions with partially dissociable functions^32,33^: the dorsal hippocampus (dHPC) contributes to precise spatial memory, the ventral hippocampus (vHPC) to more emotional and generalized representations, and the intermediate hippocampus (iHPC) has been implicated in goal-directed navigation and value-based processing^34–38^. Importantly, hippocampal cognitive maps are increasingly understood as social as well as spatial. Hippocampal circuits support social recognition and social-memory storage, including ventral CA1 and CA2–ventral CA1 pathways, and social-memory representations can be reactivated during hippocampal sharp-wave ripples^39–44^. At the population-code level, hippocampal neurons represent not only the animal’s own position, but also the position of conspecifics in rodents and bats^45–47^ — a finding whose generality is debated^48,49^, with discrepancies potentially explained by the relevance or salience of the encoded social agent —, multiplex self- and other-related variables, and encode social identity, sex, hierarchy and affiliation in group-living bats^50,51^. Particularly relevant here, Forli and Yartsev^52^ showed that hippocampal activity in freely interacting bats represents collective spatial behaviour, indicating that hippocampal maps can be structured by the movements and identities of other individuals. Together, these studies show that the hippocampus can incorporate social agents into spatial and mnemonic representations.

Consistent with this broader social role, the dorsal hippocampus is required for the expression of observational contextual fear shortly after learning^5^. Yet these literatures leave a central gap. Social-hippocampus studies show that the hippocampus can represent others as agents in space and memory, but have not determined how another individual’s affective state is bound to the observer’s spatial map. Conversely, observational-fear studies show that the hippocampus is required for recent recall, but have not tracked how hippocampal population activity evolves from witnessing another’s distress to subsequent memory expression.

We therefore recorded single-neuron activity across hippocampal subregions in male observer rats during differential contextual fear learning by observation. The task contrasted a safe context with a shock-observation context and allowed us to follow hippocampal population activity across the phases in which socially acquired threat memories are encoded, consolidated and expressed: shock observation, post-learning rest and later recall (Fig. 1a).

**Figure 1:**
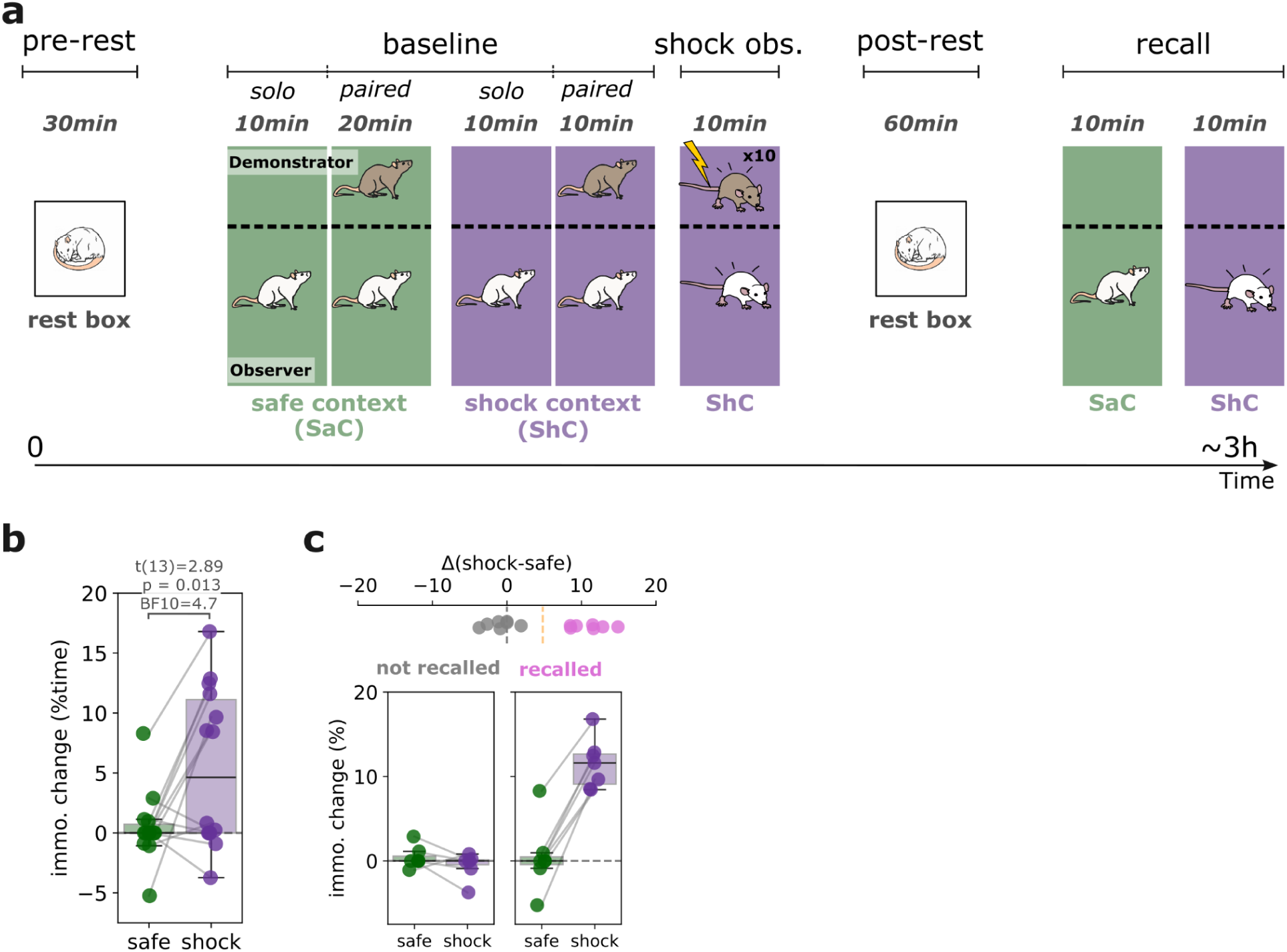
Rats recall the association between shock observation and its context. **a.** Schematic representation of the successive phases in our vicarious contextual fear learning paradigm. Colors symbolize different contexts, which differed in shading of the wall and floor plates, odor, and orientation in the Faraday cabinet. The stimulus-context association was pseudo-randomly assigned and counterbalanced across animals. The order of safe vs shock context was fixed to prevent carryover of distress from a preceding shock context to a subsequent safe context. **b.** Percentage increase in immobility as a proxy of freezing from baseline to recall phases in the safe (green) vs. shock (purple) context (N=14 rats). **c.** Same as B, but separating the half of animals without evidence of recall (left; NON-RECALLER) from those with evidence of differential recall (right; RECALLER). Boxplots center line represents the median, the box limits the 25th and 75th percentiles, and the whiskers extend to values up to 1.5 x the interquartile range; points indicate values for individual animals. In **b**, two-sided student T-test P value and Bayesian BF10 are reported. Source data are provided as a Source Data file.

## Results

### Observers express differential contextual freezing after shock observation

To determine whether rats learned to associate observed distress with a specific environment, we measured freezing behavior when the observers were later returned to the two contexts. Because the animals had experienced both contexts before shock observation, greater freezing in the shock-observation context compared with the safe context would indicate that observers formed a context-specific memory of where the demonstrator’s distress occurred. During recall, observers showed, consistent with observational contextual fear learning, a selective increase in immobility when returned to the shock context compared with the safe context (Fig. 1b; two-sided student t-test, t(13)=2.89, p=0.013, Cohen’s *d* = 0.99, 95% CI = [1.27, 8.83], BF10=4.68). Notably, individual animals varied substantially in the magnitude of this contextual freezing response (Fig. 1c). This variability was associated with distinct behavioral features (Supplementary Fig. 1); rats displaying a contextual freezing response during recall (RECALLER) showed a stronger preference for the vicinity of the demonstrator’s compartment during baseline and recall, and displayed a larger pupil response during shock observation.

### Shock observation recruits hippocampal pyramidal neurons along the septotemporal axis

We next asked whether observing another rat receive shocks recruits hippocampal neurons, and whether such responses differ along the septotemporal axis. To address this, we quantified changes in neuronal firing immediately following each shock delivered to the demonstrator using Neuropixels probes (Fig. 2a, Supplementary Fig. 2, Supplementary Table 1-2). Based on reconstructed implantation locations, we sampled dorsal, intermediate, and ventral hippocampal subregions and recorded 337 putative pyramidal neurons (73 dorsal, 144 intermediate, 120 ventral) across 14 animals (Fig. 2). Because pyramidal neurons constitute the principal substrate of the hippocampal spatial code^53^, we focused on this population, while the 149 putative interneurons we recorded are reported in Supplementary Fig. 3. Quantifying pyramidal neuron activity during the 1s following footshocks revealed shock-observation–related modulation across the hippocampus, with a significant enrichment of shock-excited neurons in the intermediate hippocampus, and shock-inhibited neurons in the ventral hippocampus (Fig. 2b- c; χ²(4)=11.83, p=0.019, Cramér’s V = 0.13).

**Figure 2:**
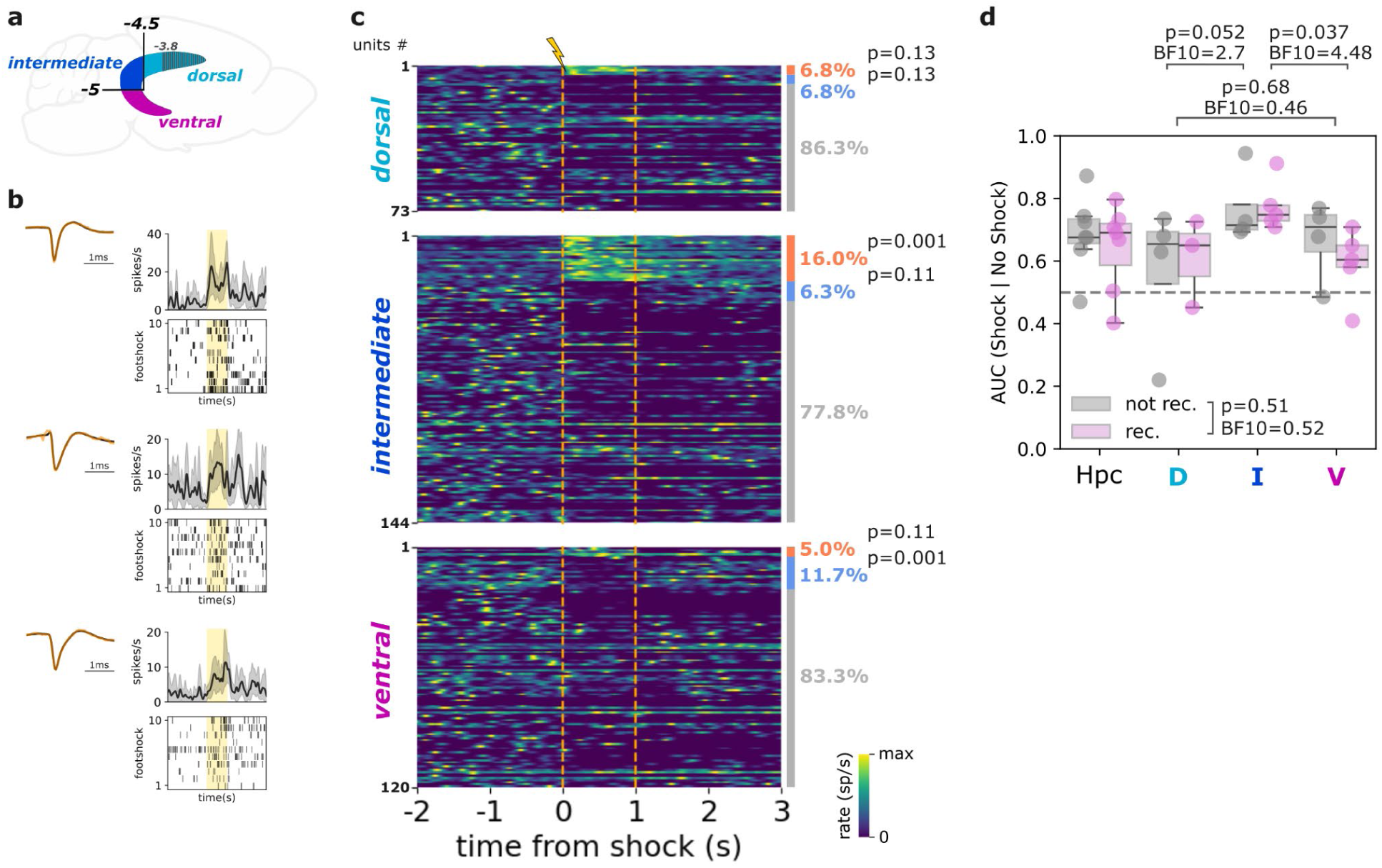
A population of putative pyramidal neurons is recruited by shock observation mostly within the intermediate hippocampus. **a.** Schematic representation of the anatomical segmentation of the dorsal (recorded between −3.8 and −4.5mm AP from bregma), intermediate (recorded posteriorly to −4.5mm and dorsally to −5mm relative to bregma), and ventral (recorded below to −5mm from bregma) hippocampus (see Supplementary Figure 1 for details). **b.** Example of three putative pyramidal neurons that increase their rate during shock observation. For each, the top right panel represents the mean firing rate with the shaded area representing the 95% confidence interval estimated from 1000 iterations of bootstrapping; bottom right, raster plot for each footshock delivery to the demonstrator; top left panel, average spike waveform for all recorded spikes throughout the whole sessions (black) or during the 10 footshocks (overlayed in orange). The shaded yellow area corresponds to the 1s of footshock delivery. **c.** Heatmap representing the mean firing rate of single putative pyramidal neurons separated by hippocampal sub-region (dorsal, intermediate, ventral). For each neuron type and sub-region, neurons are ordered from top to bottom as follows: significantly excited (orange), significantly inhibited (blue) and not significantly modulated (gray) single neurons. Overall percentages are indicated on the right side of the heatmaps. Orange dashed vertical lines represent the onset and offset of footshock delivery. **d.** Area Under the Curve (AUC) of the Receiver Operating Characteristic (ROC) curve, computed from the results of a linear classifier trained to discriminate between corrected shocks and matched control-moments without shocks from the single-units responses recorded in the entire hippocampus (Hpc, N = 14 rats) or the dorsal (D, N = 7 rats), intermediate (I, N = 9 rats) and ventral (V, N = 9 rats) subregions separately for RECALLER (rec., pink) and NON-RECALLER (not rec., grey) rats. Boxplots center line represents the median, the box limits the 25th and 75th percentiles, and the whiskers extend to values up to 1.5 x the interquartile range; points indicate values for individual animals. In **c**, two-sided Monte Carlo P value are reported; in **d**, Holm corrected P values and Bayesian BF10 from two-sided student t-test are reported. Source data are provided as a Source Data file.

We next asked whether hippocampal population activity could distinguish shock- observation moments from control moments in which the observer occupied similar locations but no shock was delivered to the demonstrator (Fig. 2d). Population activity discriminated shock- observation from matched control moments in the full hippocampal population and in the intermediate and ventral hippocampus, but not in the dorsal hippocampus (one sample t-test; for the entire population of hippocampal units: t(13) = 4.83, p = 0.00033, Cohen’s *d* = 1.29, 95% CI = [0.59 0.74], BF10>100; dHPC: t(6)=1.20, p = 0.28, Cohen’s *d* = 0.45, 95% CI = [0.41, 0.76], BF10 = 0.61; iHPC: t(8) = 8.79, p < 0.0001, Cohen’s *d* = 2.93, 95% CI = [0.7, 0.84], BF10 > 100; vHPC: t(8) = 3.15, p = 0.014, Cohen’s *d* = 1.05, 95% CI = [0.53, 0.72], BF10 = 5.01). Discriminability was strongest in the intermediate hippocampus (iHPC vs dHPC: t(14) = 2.49, p=0.052, Cohen’s *d* = 1.26, 95% CI = [0.03, 0.34], BF10 = 2.7; iHPC vs vHPC: t(16) = 2.82, p=0.037, Cohen’s *d* = 1.33, 95% CI = [0.04, 0.25], BF10 = 4.48) and did not differ between RECALLER and NON-RECALLER animals (Fig. 2d, t(12) = 0.61, p = 0.56, Cohen’s *d* = 0.32, 95% CI = [−0.11, 0.19], BF10 = 0.50). Having established that shock observation was represented at both single-neuron and population levels, we next asked whether these responses were embedded within hippocampal spatial coding.

### Shock-observation responses scale with place-field firing

A key property of hippocampal pyramidal neurons is that they fire preferentially when the animal occupies specific locations in space, forming place fields. Importantly, place cells do not only encode where the animal is: previous work has shown that non-spatial events can modulate firing within these spatial fields. During first-person auditory fear conditioning, for example, hippocampal place cells acquired responses to the conditioned stimulus that were gated by the animal’s location, such that responses occurred preferentially when the rat was within the neuron’s place field^54^. We therefore asked whether a similar spatial gating principle applies when the aversive event is not experienced directly, but observed in another rat: do hippocampal neurons respond more strongly to shock observation when the observer is near the peak of the neuron’s place field than when it is outside the field? Using place fields defined during baseline exploration of the shock context (Fig. 3a, Supplementary Fig. 4), we first observed that shock- observation responses were position dependent: individual neurons responded more when the observer was within than outside their place field. This was evident in example neurons (Fig. 3b).

**Figure 3:**
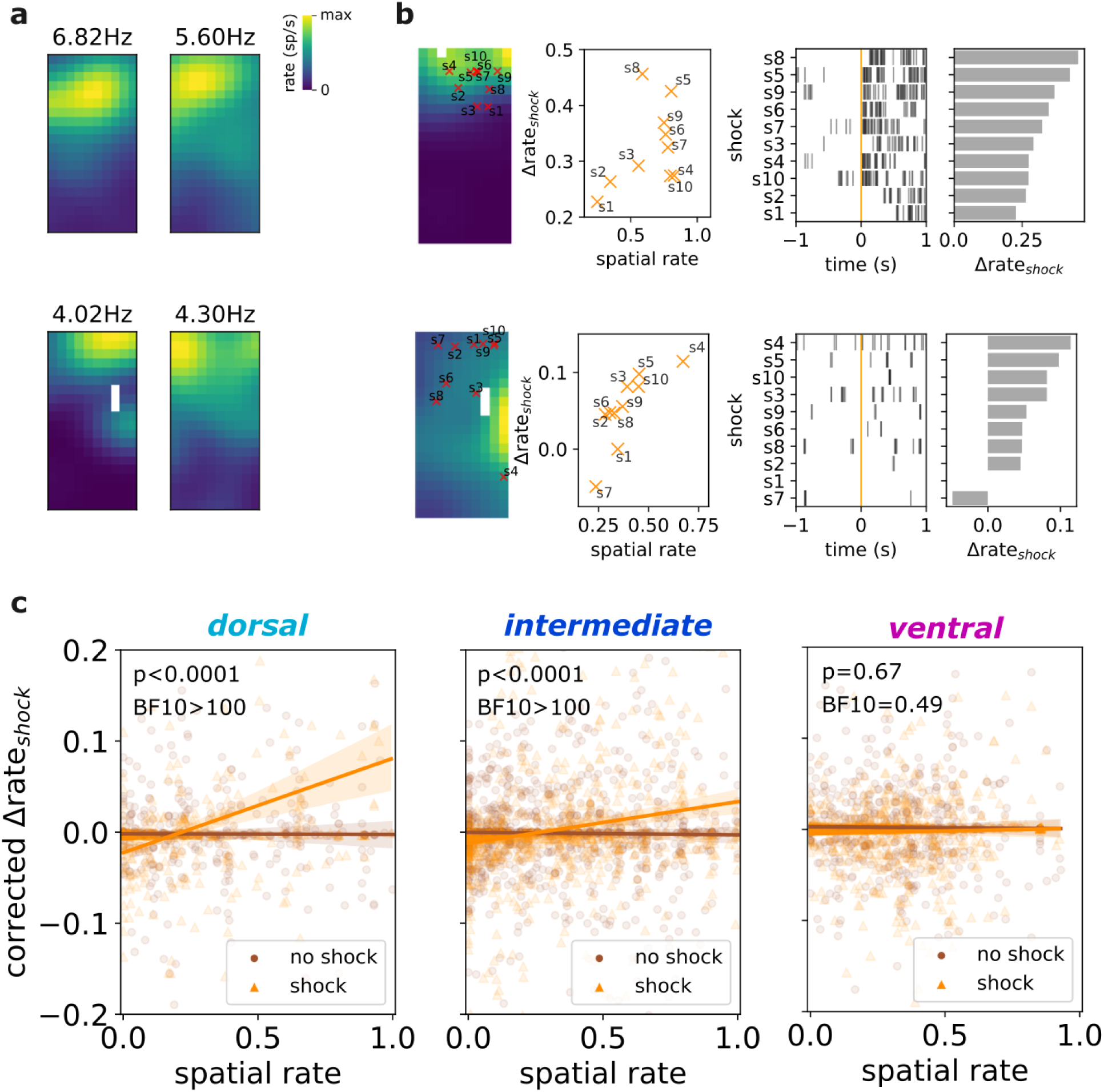
Place-field dependent modulation of shock-observation responses in the dorsal and intermediate hippocampus. **a.** Example spatial rate maps and maximum within-field rate from shock observation excited putative pyramidal cells computed from the solo baseline exploration of the observer’s compartment in the shock context. **b-c.** Modulation of the shock observation response by spatial location. **b.** Examples of two neurons, from left to right panel: position of the observer at the time of footshock delivery overlaid on single neuron’s spatial rate maps computed from the solo baseline exploration of the observer’s compartment in the shock context; raster plot of the shock observation response plotted against basal spatial firing rate; spikes raster plot relative to each footshock delivery onset and ordered by shock observation response magnitude as displayed in the last panel. **c.** For each hippocampal subregion, linear regression between corrected neuronal response for speed and proximity to the demonstrator’s compartment and estimated basal spatial firing rate at times of footshock delivery (gold) or matched control-moments without shocks (sienna). In **c**, mean regression line with 95% confidence interval. Dots represent single trial responses. Uncorrected two-sided P value and BF10 for the linear mixed model’s fixed effect of the interaction: baseline spatial rate X footshock delivery times. Source data are provided as a Source Data file.

To test this systematically across the population, we treated each shock as a separate trial and asked whether the magnitude of each neuron’s shock-observation response depended on the neuron’s baseline spatial firing rate at the observer’s position during that trial. This baseline spatial firing rate provides a continuous measure of how close the observer was to the neuron’s preferred firing location, from out-of-field positions to the place-field peak. This analysis required controlling for two potential confounds. First, hippocampal firing can be modulated by movement speed^55^. Second, because observers spent much of the shock-observation period near the divider, and because shock-responsive neurons could be spatially biased toward that region, an apparent place × shock interaction could arise if neurons simply responded more strongly when the animal was close to the demonstrator. We therefore corrected shock-observation responses for speed and proximity to the demonstrator’s compartment and compared shock trials with spatially matched control moments without shocks. Under a conjunctive coding account, the relationship between baseline spatial firing and neuronal response should be stronger during shock observation than during spatially-matched control moments.

Consistent with this prediction, shock-response magnitude scaled with baseline spatial firing rate at the observer’s position after controlling for speed and proximity to the demonstrator (Fig. 3c). Linear mixed-model analyses revealed a robust interaction between spatial firing and shock observation in the dorsal and intermediate hippocampus (dHPC: β = 0.091 [0.068, 0.12], Z = 7.55, p < 0.0001, BF₁₀ > 100; iHPC: β = 0.043 [0.026, 0.059], Z = 5.09, p < 0.0001, BF₁₀ > 100), but not in the ventral hippocampus (β = 0.004 [−0.014, 0.022], Z = 0.43, p = 0.67, BF₁₀ = 0.49) and with the absence of difference between rats displaying contextual freezing during recall versus not (dHPC: β = 0.01 [−0.03, 0.06], Z = 0.58, p = 0.57, BF₁₀ = 0.13; iHPC: β = 0.00 [−0.033, 0.033], Z = 0.004, p = 0.99, BF₁₀ = 0.62). These results identify a gain-like conjunctive coding mechanism by which hippocampal neurons encode where the observer was when witnessing the others’ distress. Consistent with the proposed role of the intermediate hippocampus in integrating spatial and value-related information^34–38^, shock-observation–responsive neurons were more prevalent in this region.

### Shock-context spatial representations are selectively stabilized in the intermediate hippocampus

The hippocampal spatial code is not static: learning induces reorganizations and stabilization of place-cell representations depending on the behavioral significance of an environment^20,21,56^. We therefore asked whether observing shocks altered the spatial representations associated with the safe and shock contexts, and whether these changes differed across hippocampal subregions. We compared place-field representations between solo baseline and recall. To quantify how much spatial representations changed across learning, we compared each neuron’s place field between baseline and recall using correlation distance, defined as 1-r between spatial firing maps. We refer to this measure as baseline-to-recall correlation distance: higher values indicate greater map reorganization, whereas lower values indicate greater map stability. While neurons showed no systematic changes in excitability or spatial information content (Supplementary Fig. 4b), baseline-to-recall correlation distance was substantial across all hippocampal subregions in both the safe and shock contexts (Fig. 4b). Notably, in the intermediate hippocampus, baseline-to-recall correlation distance was lower for the shock context than for the safe context, indicating greater stability of shock-associated spatial representations (iHPC: β = −0.22 [−0.37, −0.077], Z = −3.01, p = 0.003, BF₁₀ = 18.01), an effect not observed in dorsal or ventral hippocampus (Fig. 4b). Shock-observation–responsive pyramidal neurons in the intermediate hippocampus showed lower baseline-to-recall correlation distance than nonresponsive neurons (β = −0.28 [−0.775, −0.018], Z=-2.1, p=0.036, BF10=2.99), providing suggestive evidence that shock-responsive cells preferentially stabilized their spatial tuning.

**Figure 4:**
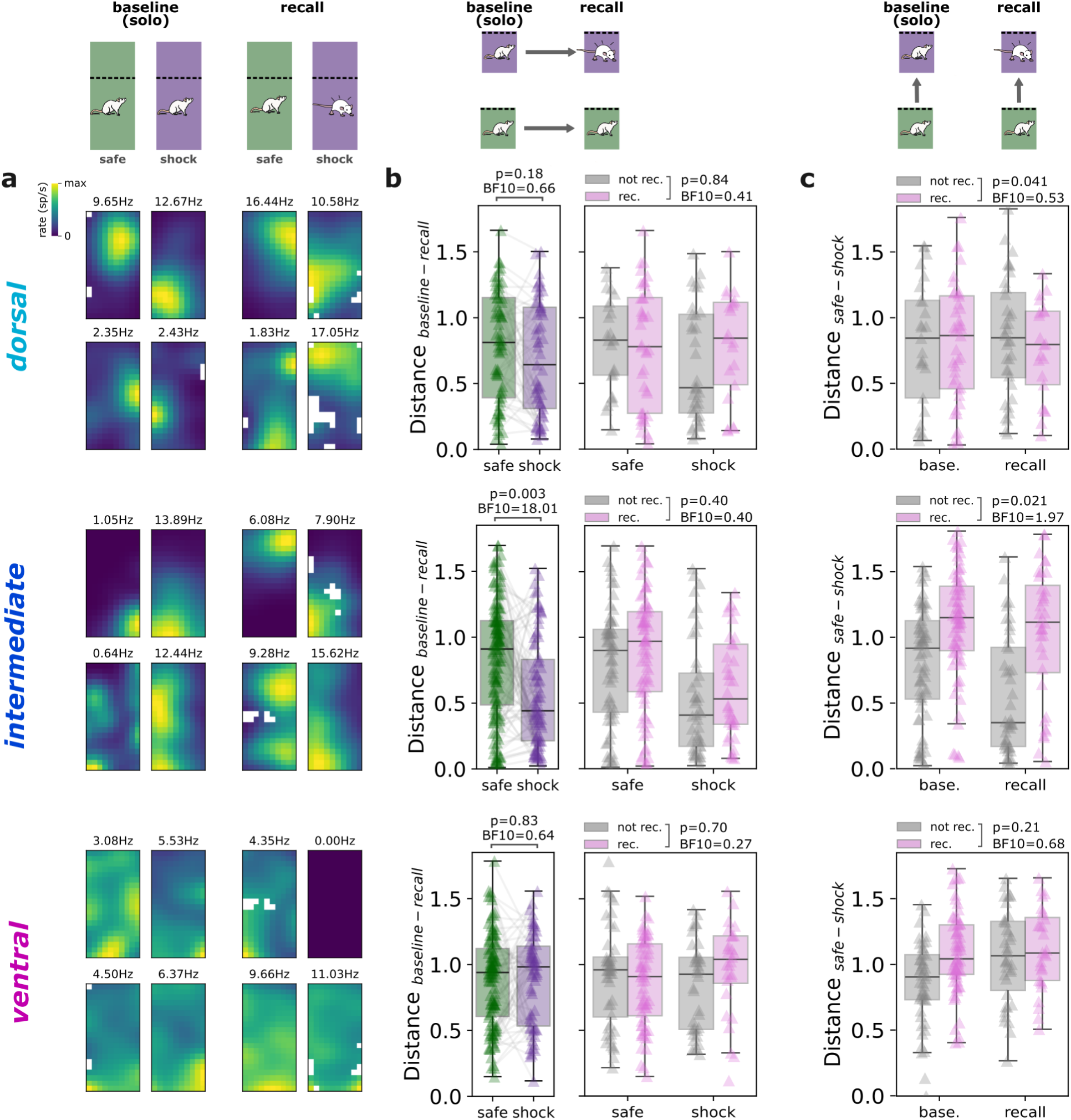
Stabilization of the spatial representation associated with the shock observation in the intermediate hippocampus. Row corresponds to neurons recorded from a different subregion of the hippocampus**. a.** Example spatial rate maps from putative pyramidal neurons during the solo exploration of the observer’s compartment in the safe and shock context during baseline and recall, and corresponding peak field rate. **b.** Baseline-to-recall correlation distance (1-r) between spatial firing maps, shown separately for safe (green) and shock (purple) contexts. Lower values indicate greater map stability across learning. Left panels show all animals; right panels show the same analysis further split for RECALLER (rec.) or NON-RECALLER (not rec.) observer rats; for neurons recorded from the dorsal (N = 73 neurons, 34/39 from RECALLERS/NON-RECALLERS), intermediate (N = 144 neurons, 78/66 from RECALLERS/NON-RECALLERS), and ventral (N = 120 neurons, 66/54 from RECALLERS/NON-RECALLERS), **c.** Safe–shock correlation distance (1-r) between spatial firing maps for the same neurons, shown separately for baseline and recall. Higher values indicate greater separation between the spatial representations of the two contexts. Data are further split for RECALLER (rec.) or NON-RECALLER (not rec.) rats. Boxplots center line represents the median, the box limits the 25th and 75th percentiles, and the whiskers extend to values up to 1.5 x the interquartile range; points indicate values for individual neurons. Uncorrected two-sided P values and BF10 for fixed effects of linear mixed models. Source data are provided as a Source Data file.

Because only a subset of animals later expressed differential contextual freezing, we next asked whether hippocampal spatial maps differed between RECALLER and NON-RECALLER animals in either stability or context separation. For stability, we again used baseline-to-recall correlation distance. For context separation, we computed safe–shock correlation distance, defined as 1-r between spatial maps from the safe and shock contexts, where higher values indicate more distinct representations of the two contexts. We found no difference in shock-context baseline-to-recall correlation distance between RECALLER and NON-RECALLER rats (Fig. 4b; β = 0.083 [−0.11, 0.28], Z=0.82, p = 0.40, BF10=0.40). However, intermediate hippocampal place cells showed greater safe–shock correlation distance during recall in RECALLER animals (Fig. 4c; β = 0.26 [0.038, 0.47], Z = 2.30, p = 0.021, BF10=1.97), consistent with their stronger behavioral discrimination between the two contexts.

### Post-learning SWRs preferentially reactivate shock-context ensemble patterns

Hippocampal activity patterns expressed during experience are often reactivated during subsequent sharp-wave ripple (SWR) events occurring during rest or sleep. Such reactivations contribute to memory consolidation^26,28,29^ and the stabilization of spatial representations^57^ by preferentially reinstating patterns of coordinated neuronal activity associated with behaviorally relevant salient experiences^58,59^. We therefore asked whether ensemble activity patterns expressed during exploration of the shock context were preferentially reactivated during post-learning rest. To address this question, we compared the pattern of pairwise correlations between simultaneously recorded neurons during exploration across the entire baseline phase, in both solo and duo conditions, and during post-learning SWRs. Conceptually, this analysis asks whether neurons that tended to co-fire together during exploration also tended to co-fire together again during subsequent rest. If shock-context representations are preferentially consolidated, then the structure of neuronal coactivity during SWRs should resemble that observed during exploration of the shock context more strongly than that of the safe context. Importantly, some neurons may exhibit stable correlations across all recording periods independently of learning. To account for these pre-existing patterns, we used an explained-variance^60^ analysis that statistically removes the similarity already present before shock observation. Forward explained variance quantifies how strongly post-learning SWR activity resembles exploration-related activity after controlling for pre-learning rest. As a control analysis, we also computed reversed explained variance, which asks whether pre-learning rest resembles exploration after accounting for post-learning rest. Preferential reactivation associated with learning is therefore expected to produce higher forward than reversed explained variance (Fig. 5 and Supplementary Fig. 5). Reactivation did not differ between forward and reversed explained variance for the safe context (t(13) = 0.087, p = 0.93, Cohen’s *d* = 0.033, 95% CI = [−0.11, 0.12], BF10 = 0.27), indicating no neuronal activity pattern reactivation of that safe context. In contrast, for the shock context, forward explained variance robustly exceeded reversed explained variance (t(13) = 6.56, p < 0.0001, Cohen’s *d* = 2.08, 95% CI = [0.17, 0.33], BF10 > 100; Fig. 5b), indicating that post-learning SWRs preferentially reinstated ensemble coactivity patterns expressed in the shock context. To determine whether this reactivation reflected the shock context broadly or the shock-observation moments specifically, we repeated the analysis using only activity from the shock-observation phase. Forward explained variance was again elevated, but did not differ from that obtained using the entire shock-context exploration period (t(13) = 1.33, p = 0.21, Cohen’s *d* = 0.13, 95% CI = [−0.01, 0.06], BF10 = 0.56). Thus, post-learning reactivation was not selectively enriched for coactivity patterns unique to shock-delivery moments, but instead primarily reflected the broader population structure associated with the shock context.

**Figure 5:**
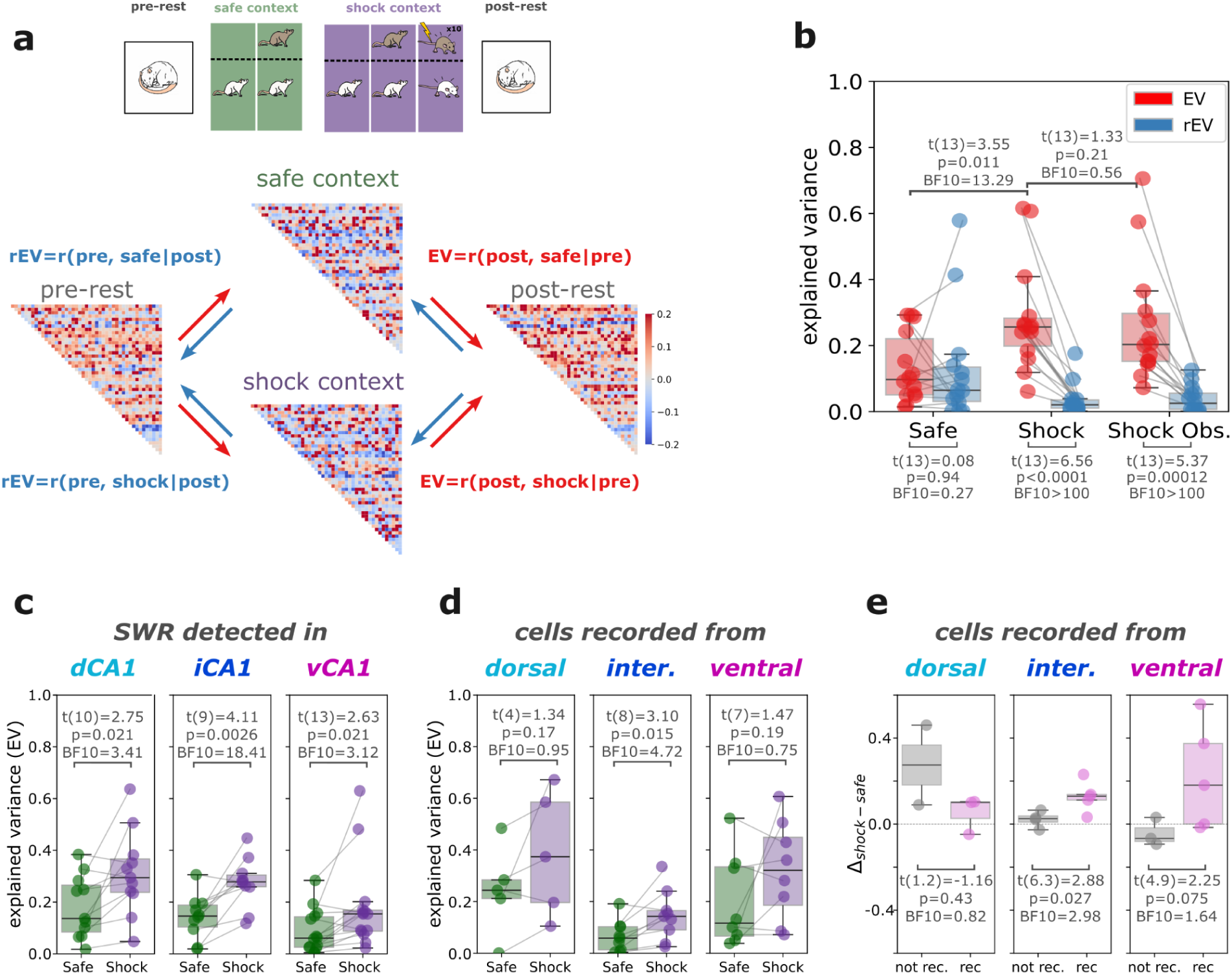
Enhanced reactivation of the patterns of activity associated with the shock context within the intermediate hippocampus. **a.** Schematic representations of the statistical approach employed to compute forward and explained variance separately for patterns of activity associated with the exploration of the safe and shock context, together with example cell-pairs correlation matrices from one animal. **b.** Forward (red) and reverse (blue) explained variance for spike associated with the entire exploration duration within the safe and shock context, as well as restricted to the shock observation phase (N = 14 rats). **c.** Forward explained variance for spike associated with the entire exploration duration within the safe (green) and shock (purple) context and restricted to neuronal activity co-occurring with SWR detected in the dorsal (dCA1, N=11 rats), intermediate (iCA1, N=10 rats) and ventral (vCA1, N=14 rats) hippocampal subregions. **d.** Forward explained variance for safe and shock contexts, separated by the anatomical location of recorded neuron pairs (dorsal: N = 5 rats, intermediate: N = 9 rats, ventral: 8 rats). **e.** Difference in forward explained variance between shock and safe contexts, separated by recall status (rec, in pink: RECALLER, not rec, in grey: NON-RECALLER; dorsal: N = 3/2 RECALLER/NON-RECALLER rats, intermediate: N = 5/4 RECALLER/NON-RECALLER rats, ventral: 5/3 RECALLER/NON-RECALLER rats). Boxplots center line represents the median, the box limits the 25th and 75th percentiles, and the whiskers extend to values up to 1.5 x the interquartile range; points indicate values for individual animals. Holm corrected P values and BF10 from two-sided student t-tests. Source data are provided as a Source Data file.

We next asked whether this effect could be explained by global changes in SWR activity from pre- to post-learning rest. This was not the case: post-learning rest did not show a systematic increase in SWR rate, within-SWR firing rate, or mean cell-pair correlation that could account for the preferential reactivation of shock-context ensemble patterns (Supplementary Fig. 5d-f). Having established that post-learning SWRs preferentially reinstated shock-context activity patterns, we then asked where along the hippocampal axis this effect was most pronounced. Reactivation was stronger for the shock than the safe context (t(13) = 3.55, p = 0.029, Cohen’s *d* = 1.12, 95% CI = [0.06, 0.25], BF₁₀ = 13.29), and this bias was observed for SWRs detected in dorsal, intermediate and ventral hippocampus (Fig. 5c). However, the effect size was largest for SWRs detected in the intermediate hippocampus. Consistent with this, when analyses were separated by the anatomical location of recorded neuron pairs, a significant shock–safe difference in forward explained variance was confined to intermediate hippocampal neuron pairs (Fig. 5d; t(8) = 3.10, P = 0.015, Cohen’s *d* = 1.00, 95% CI = [0.02, 0.14], BF₁₀ = 4.72).

Finally, we asked whether preferential shock-context reactivation was related to later behavioral expression of contextual fear. The shock–safe difference in forward explained variance among intermediate hippocampal neuron pairs was larger in RECALLER than NON-RECALLER rats (Fig. 5e; t(6) = 2.88, P = 0.026, Cohen’s *d* = 1.80, 95% CI = [0.02, 0.2], BF₁₀ = 2.98). Together, these results indicate that post-learning SWRs preferentially reinstated ensemble activity patterns associated with the shock context, with a dominant contribution from the intermediate hippocampus and stronger reactivation in animals that later expressed contextual freezing.

## Discussion

### Summary

Using an adapted contextual fear learning-by-observation paradigm, we investigated how witnessing a conspecific in distress reshapes hippocampal representations. Observer rats were presented to two previously neutral contexts (safe and shock) alone, then in the presence of a demonstrator, and witnessed the demonstrator receive electrical footshocks while in the shock context. During later recall, observers displayed increased freezing selectively in the shock context at the group level, although with marked inter-individual variability. Animals that later discriminated between safe and shock contexts showed stronger pupil dilation during shock observation and greater proximity to the demonstrator.

Single-unit recordings from the hippocampus suggest a prominent role for the intermediate hippocampus in socially acquired contextual threat. First, shock observation elicited a spatial-firing-rate-dependent excitatory response in pyramidal neurons, particularly in the intermediate hippocampus, where a larger proportion of neurons encoded shock observation and where population activity more accurately decoded shock-related moments. Second, neuronal activity patterns associated with the shock context were preferentially reactivated during subsequent rest sharp-wave ripples, again most prominently in the intermediate hippocampus. Third, contextual representations underwent widespread remapping following shock observation, with a relative local stabilization of the shock-context spatial representations within the intermediate hippocampus.

### Place-Shock coding in the dorsal and intermediate hippocampus

A central finding of this study is that in both dorsal and intermediate hippocampus, pyramidal neurons exhibited place-field-dependent modulation of their firing rate during shock observation, providing a potential neuronal substrate to integrate socially conveyed threat with spatial information. These findings place the hippocampus within the broader network of structures (anterior cingulate cortex, insula, medial prefrontal cortex, amygdala)^6,7,61–63^ in the mammalian brain, displaying single neuron responses to socially observed aversive experiences. They expand on the accumulating evidence for the integration of relevant social stimuli within the hippocampal population code, such as conspecific attributes^39–43,51^ and location^45–47^, and a more global reorganization of the hippocampal spatial code by social group dynamics^47,50–52^. Previous work has shown that hippocampal place cells in the dorsal hippocampus can multiplex spatial and non-spatial variables through firing-rate modulation, including task rules, sensory cues and first-hand aversive experiences^10–12,54^. The present findings extend this framework to vicarious aversive experience and to the intermediate hippocampus.

Reward is also known to modulate both the dorsal and intermediate hippocampus spatial representations^36,44,56,64,65^, and hippocampal neurons are sensitive to salience, attention and behavioral state. In our task, pupil dilation during shock observation is consistent with heightened arousal or attention, whereas proximity to the demonstrator’s compartment indicates active social monitoring^66,67^. Together, these observations raise the possibility that the observed hippocampal responses reflect a socially derived salience- or value-place representation linked to the demonstrator’s distress and the observer’s affective response to it^34–38^.

### Widespread remapping associated with the specific stabilization of shock observation responsive neurons spatial maps

In the present study, we found a widespread reorganization of the spatial representations of both the shock and safe context following shock observation across all hippocampal subregions. The reorganization of hippocampal cells spatial firing is thought to reflect representational changes^16,17,20,68,69^ and their stabilization has been associated with memory processes^70–72^. The changes in spatial firing associated with the shock context are thus aligned with this model and with previous work investigating first-hand fear learning^20,68,69,72^, suggesting vicarious learning induced changes in the associated spatial representations. The larger changes associated with the safe context are however surprising relative to previous report^69^ and the reported stability of the hippocampal code in unchanging conditions^73–75^, and they may have reflected interaction between the spatial representations of the two contexts and/or active safety learning^68,76^, emphasized by the relatively subtle sensory dissociations between the two contexts. Our experimental design prevents from completely ruling out an effect of time, although it is unlikely in the training conditions and time scale used^73–75^.

Despite this global remapping, we also found a relative stabilization of the spatial representations of the shock context for neurons within the intermediate hippocampus and this stabilization was most markedly true for neurons that were excited by shock observation. Enhanced arousal or attention during shock observation may have contributed to this stabilization^21,66,77^, potentially through neuromodulatory influences such as locus coeruleus- dependent plasticity mechanisms^78–80^. This dissociation between widespread reorganization of spatial representations associated with the stabilization of the spatial firing of neurons to aversive events could be a mechanism through which subjective value is anchored to the updated spatial representations supporting episodic memory in a familiar environment.

Interestingly, the stabilization of shock-context related spatial representations did not differ between animals displaying different freezing behavior during recall. Rather, animals displaying differential contextual freezing also showed stronger neuronal representational distance of the two contexts. Although anecdotal, these observations are consistent with the hypothesis that NON- RECALLER animals failed to maintain sufficiently distinct representations of the safe and shock contexts^81–83^. Because these animals showed little freezing in either context, their behavior is better interpreted as safety-dominated contextual ambiguity than as fear generalization: The shock-observation experience may have been encoded against a broader familiar-arena representation, established during habituation and baseline exploration, so that threat-related information in the shock context was diluted by overlapping safety-related representations^76,84,85^. Such dilution could prevent robust contextual freezing despite encoding of the shock-observation episode.

### Preferential reactivation of shock-context activity patterns during post-learning sharp-wave ripples

During post-observation rest, neuronal co-firing patterns^31,60,86^ associated with the shock context were preferentially reactivated during sharp-wave ripple (SWR) activity compared to patterns associated with the safe context. Notably, neuronal activity patterns associated with the shock observation moments were no more preferentially reactivated than those associated with the overall active experience within the associated context. Preferential reactivation or replay of salient, valuable and emotional experience^38,58,59,87–90^ has been observed previously; the present findings extend this principle to vicarious aversive experiences.

We found enriched shock-related neuronal reactivation associated with SWR activity detected in all three hippocampal subregions: dorsal, intermediate and ventral. These results suggest that activity in all subregions in the hippocampus contributed to the preferential reactivation of shock-context-related content. Although, we found some indication for enriched cross-region SWR occurrence with short <20ms delays, whether such reactivations occurred locally or synchronously within the hippocampal subregions^91^ such as reported between dorsal and ventral hippocampus for first-hand aversive experiences^92^ will require further investigation.

Consistent with the enhanced recruitment of neurons within the intermediate hippocampus during shock observation, we found that the preferential reactivation of the shock context related content was strongest for SWR activity and for neurons recorded within the intermediate hippocampus consistent with previous work reporting reactivations for spatial representations associated with high value in this structure^38^. The preferential reactivation of shock-context activity in the intermediate hippocampus may reflect two complementary processes. First, stronger neuronal responses during shock observation may have induced synaptic plasticity, biasing subsequent SWRs toward the ensembles active in the shock context^93^. Second, such post-learning reactivation may itself contribute to memory consolidation^26,28,29,58^, potentially by stabilizing shock-context spatial representations^57^ in intermediate hippocampal place cells. Consistent with this latter possibility, shock–safe differential reactivation was stronger in rats that later showed shock-context-specific freezing.

### Scope and Limitations

Several features of the present design define the scope of these conclusions. First, although several hippocampal signatures were related to individual differences in recall, not all components of the proposed mechanism distinguished animals that later did or did not express differential freezing. RECALLER animals showed greater separation between safe- and shock-context spatial maps during recall in the intermediate hippocampus, and stronger shock–safe differential reactivation in intermediate hippocampal neuron pairs during post-learning SWRs. These findings suggest that the behavioral expression of contextual freezing may be associated with how distinctly the two contexts were represented at recall and how strongly shock-context ensemble patterns were reinstated during post-learning rest.

However, other neural phenomena were present at the group level without reliably separating RECALLER from NON-RECALLER animals. In particular, observation-phase shock responses, place × shock conjunctive coding and shock-control population discriminability were robustly detected, especially in the intermediate hippocampus, but did not predict later differential freezing. Similarly, stabilization of shock-context spatial representations was strongest in the intermediate hippocampus but did not differ reliably between RECALLER and NON-RECALLER animals. These observations do not rule out a role for these neural responses in memory encoding. Rather, they suggest that these responses are not sufficient on their own to predict the behavioral expression of contextual freezing. An animal may encode the shock-observation experience without expressing differential freezing at recall, either because safety signals suppress freezing in that context, because the memory generalizes across contexts leading to safety learning associated to the arena, or because downstream circuits fail to translate the memory into freezing behavior. Causal manipulations of hippocampal activity during observation, consolidation or recall will be required to determine how these representations contribute to memory formation and behavioral expression.

Second, observer rats were pre-exposed to footshocks before observational learning. This manipulation was used to promote robust observational freezing^2,94^, but it means that the hippocampal mechanisms described here may reflect the integration of witnessed distress with prior first-hand aversive experience. This is important because prior shock experience can alter the circuits supporting observational fear^95^, and future experiments should test whether similar place–shock conjunctive coding, spatial stabilization and post-learning reactivation emerge in fully naïve observers.

Third, our conclusions about regional specificity should be interpreted in light of the anatomical distribution of recordings. Although we sampled dorsal, intermediate and ventral hippocampal subregions, sampling was not uniform along the septotemporal axis, and the intermediate hippocampus contributed the largest pyramidal-cell sample. The present data therefore provide strongest evidence for a mechanism in the intermediate hippocampus, where shock-observation responses, spatial stabilization and post-learning reactivation converged. By contrast, weaker or non-significant effects in dorsal and ventral hippocampus should not be interpreted as evidence that such mechanisms are absent from those regions, particularly given the more limited coverage of the full septotemporal axis, including the most anterior/septal dorsal hippocampus. Additionally, SWR detection relied on a fixed ripple-frequency band (150–250 Hz) across subregions. Given reports that ripple peak frequency decreases along the dorsoventral hippocampal axis^101,102^, this approach may have been comparatively conservative for detecting ripples in the ventral hippocampus, potentially underestimating reactivation effects in this region. Fourth, recordings were performed in male rats. Although emotional contagion is behaviorally similar across male and female rats^96^, the generality of our hippocampal findings to females remains to be established. Future studies combining denser longitudinal sampling, both sexes, fully naïve observers and region-specific causal manipulations will be needed to determine how broadly these mechanisms extend across the hippocampal axis and whether intermediate hippocampal activity is necessary for observational contextual fear memory.

## Conclusion

Together, our findings identify a candidate neuronal mechanism through which observed distress becomes situated within the hippocampal cognitive map. Although it was known that animals freeze when returned to a context in which they witnessed shocks^5,7,97^, the neurocomputational principles by which such vicarious threat information is embedded in space were unknown. We show that for hippocampal neurons, particularly in the intermediate subregion, shock-observation responses scale continuously with spatial firing rate, forming a graded population code that maps others’ distress onto the observer’s spatial position. This spatially modulated representation is selectively stabilized and preferentially reactivated, consistent with a mechanism by which socially acquired threat information becomes anchored within the hippocampal map of space.

## Methods

### Animal model

Twenty-eight 6-8 week-old male Long-Evans rats were obtained from Janvier, France. Each animal was randomly assigned to either the observer (n=14) or the demonstrator (n=14) group. Upon arrival, all animals were housed in groups of 4 (2 demonstrators and 2 observers) in open-topped type IV cages. Following surgical implantation, the observers were solitary housed in open-topped type III cages that were placed right next to the demonstrator’s home cage. The animals were housed in cages with corn cob bedding, enriched with wooden blocks and nesting shredded cartons. They were maintained on a reversed 12:12 light:dark cycle at ambient room temperature (22-24°C, 55% relative humidity, SPF) and with food pellets and water provided *ad libitum*.

All experimental procedures were performed under Licence AVD 80100 2020 9724 provided by the Centrale Commissie Dierproeven of the Netherlands and after the positive advice from the welfare body of the Netherlands Institute for Neuroscience (IVD) under project number NIN223704.

### Experimental procedure

#### Behavioral procedure

Following arrival, the animals were left untouched in their homecages for 5 to 7 days to acclimate and were next handled in the stable daily by the experimenter for 5 days. The observer animals then underwent surgical implantation and were given 3-5days of postsurgical recovery period based on the animal’s recovery and welfare assessment.

Two days prior to the experimental day, the observer animals were habituated to the sleeping box and the observer’s compartment in the experimental arena for 5min alone and 5min with the demonstrator present in its compartment. On the first day of habituation, and at least 3h after the habituation phase, the observers were shock pre-exposed in a different room as previously described^6^: after 5min of baseline, the animals received 4 electrical footshocks (0.8mA for 1s) with inter-shock intervals ranging from 240-360s. The observer was then left in its homecage in the same room to calm down for a period of at least 30min before being brought back to the stable. On the second day of habituation animals were through the same as in day 1, without the shock experience.

The experimental day (Fig.1A) was initiated with a first 30min rest phase in the sleeping box. The observers were then allowed to freely explore their compartments alone (10min) and then with the demonstrator present in its dedicated compartment (20min), first in the safe context and then in the shock context. Within the last 10min in the shock context, 10 electrical footshocks (1s at 1.5mA) were delivered to the demonstrator with pseudo-random inter-shock intervals ranging from 45 to 75s. In addition, following each footshocks, 10 control shocks were delivered at random times to an empty grid located approximately 5cm under the observer’s compartment. The context presentation order was fixed to prevent the emotional states induced by the shock delivery, for both observer and demonstrator rats, from being propagated to the safe context. Immediately following the shock observation, the observer animals were placed back in the sleeping box for 1h of rest, and finally they were brought back to the observer’s compartment alone for recall tests, first in the safe and then in the shock context.

#### Apparatus

Habituation and contextual social-fear paradigm were conducted in a closed Faraday cabinet and in dim light conditions. The experimental arena consisted of a two-compartment plexiglass box (L:75cm, W:25cm, H: 60cm). The demonstrator’s compartments had a surface area of 25x25cm with a custom-made stainless steel grid floor. The observer’s compartment occupied the remaining box area (25x50cm) with a perforated plexiglass floor. Both compartments were separated by a transparent divider also perforated on the bottom 5cm to give access to auditory, visual, and olfactory social cues. As a control, a copy of the demonstrator’s stainless steel grid floor was placed approximately 5cm underneath the observer’s ground floor near the divider. During the shock observation phase, the demonstrator’s compartment was closed with a transparent plexiglass plate to prevent the animal from jumping out of the arena.

On the experimental day, the shock and safe contexts were differentiated using different shadings of the wall and floor plates, each also odorized with vanilla and almond scents, and a 180-degree rotation of the experimental arena within the cabinet. The stimulus-context association was pseudo-randomly assigned and counterbalanced across animals. During the 2-3min necessary to apply the context changes, observer animals were placed in the rest box (see below).

During training, behavior and ultrasonic vocalization were recorded using an overhead camera (30fps, Basler acA1300) and a condenser ultrasound microphone (Avisoft-bioacustics, CM16/CMPA, Germany) placed at the top of the experimental arena. Electrical shocks were delivered via a shocker box (ENV-414, Med Associates, Inc).

During habituation and rest, observers were placed in a black plastic conical sleeping box ( ø: 15cm, H: 50 cm) located in the training room, next to the experimental cabinet, with bedding material covering the floor and food pellets provided *ad libitum*. The observers’ behavior was monitored with an overhead camera (30fps, Basler acA1300).

To avoid contextual fear generalization, footshocks pre-exposure was performed in a different room, in a rectangular transparent plexiglass box (L: 20cm, W: 15cm, H: 40cm), in a soundproof cabinet, and in red light. To further differentiate the pre-exposure context from the experimental arena, the box was washed with citrus-scented soap. The electrical shocks were delivered via a shocker box (ENV-414, Med Associates, Inc) and behavior was monitored with a side camera (Basler acA1300).

#### Surgical procedure

Animals were first injected subcutaneously with meloxicam (2mg/kg) and butorphanol (2mg/kg) 30min before the surgical intervention proper. Throughout the surgery, the animals were maintained under anesthesia with isoflurane gas, at 4-5% for the induction and between 1-3% through a mask thereafter. Their eyes were protected with a moisturizer and covered, and their body temperature was maintained at 37 °C with a heating pad.

The stereotaxic implantation of the Neuropixels implant occurred similarly to what was described previously^98^. The hair on the animal’s head was trimmed and the head aligned onto a stereotaxic frame. The scalp was cleaned with an antiseptic solution (hibicet). An incision along the midline was performed with a scalpel to expose the skull, and alm retractor were used to maintain the skin open. The surface of the skull was then cleaned out of biological material and roughened with a bone elevator. Screws were placed in a semi-circle around the future position of the implants, one of the screws, located above the cerebellum, was connected to the implant and served as a ground. One craniotomy above the right hemisphere -or two bilaterally for dual- Neuropixels implantations- were centered at −4.5mm AP and 3.2mm ML from Bregma. Small incisions were made in the dura to allow for the probes insertions. Neuropixels shanks were dipped into lipophilic dye (BioTracker 655 RED SCT108, Mercks) and slowly lowered inside the brain 7to8mm from the cortical surface. The implant was then secured in place with Metabond (C&B Metabond R&D) covered with a layer of light-curable dental cement. Finally, the skin was sutured so as to overlap with the exposed dental cement. For 10 animals, a circular camera ring was screwed to the base of the Neuropixels implant to enable anchoring the facial camera during experiment.

Immediately following surgical implantation, the animals received subcutaneous injections of 2mL of sterile saline solution and of metacam (2mg/kg) to ease recovery and hydration. The next day, and for the following three days, carpofen was administered to the animals via drinking water (0.09mg/mL).

#### Electrophysiological recordings

A total of 14 animals were successfully implanted with a chronic custom-made Neuropixels implant^98^, 5 of which carried a single Neuropixels probe 1.0 (Imec), and the remaining 9 carried two 4-shanks Neuropixels probe 2.0 (Imec). During recordings the animals were tethered to a Neuropixels recording system (PXIe, National Instrument). Raw data were acquired at 30KHz using SpikeGLX software on a dedicated computer. Behavioural events were registered with TTL recorded with spikeGLX via a digital input-output board (National Instruments) connected to the Neuropixels PXIe.

#### Head-mounted camera recordings

For 2 of the single-probe and 8 of the dual-probe implants, an additional custom-made head-mounted camera (50fps) was anchored on a dedicated rig to record the animal’s face for the duration of the experiment. For those animals during recordings, an additional tether connected the camera to a USB grabber, which in turn was connected to the acquisition computer.

#### Histological identification of implantation sites

Animals were deeply anesthetized with an IP injection of pentobarbital (butorphanol, 60mg/kg) while under isoflurane gas anesthesia (4%). The brain tissue was fixated via a transcardiac perfusion of 0.1M phosphate-buffered saline (PBS) solution, followed by paraformaldehyde solution (PFA 4%). The brains were then extracted and placed in PFA (4%) for post-fixation. The brains were sectioned in 50µm thick coronal slices with a vibratome (VT1000S, Leica). The sections from the dual-Neuropixels implantations were further processed for immunohistological staining to highlight area CA2 within the hippocampus.

Sections were first incubated in blocking solution containing (10% normal donkey serum (NDS, Sigma-Aldrich: S30-M) and 0.5% Triton-X (Sigma-Aldrich, CAS Number: 9036-19-5) in 0.1 M PBS) for 30 min at room temperature. The samples were then incubated overnight in a first bath of primary antibody (anti-PCP4, rabbit; Prestige Antibodies, Cat. No.: HPA005792, Lot.: 000040758, RRID: AB_1855086; 1:100) and after thorough washing with 0.1M of PBS, they were incubated in a secondary antibody solution (Alexa Fluor® 488 AffiniPure™ Donkey Anti-Rabbit IgG (H+L); Jackson Immunoresearch, Cat. No.: 711-545-152, Lot.: 169687, RRID: AB_2313584; 1:500) for 2 hours. All brain slices were further washed with PBS and mounted on glass microscopy slides with DAPI-containing mounting medium (4’,6-Diamidino-2-phenylindole dihydrochloride, Vectashield, Sigma-Aldrich). Afterward, all brain slices were imaged with an Axio Scan Z1 fluorescent slide scanning microscope (Zeiss) at 10x magnification (Plan-Apochromat 10x/0.45 M27). Images were acquired using Zeiss Zen software (3.7), with a pixel resolution of 0.652µm x 0.652µm and 16-bit depth with a digital CMOS camera (ORCA-Flash4 V3; Hamamatsu Photonics K.K.). Post-acquisition processing was performed in QuPath (v0.2.3), for individual thresholding of each color channel applied to the entire images.

The anatomical classification of the hippocampal subregions was done according to previous work^37,99^. All recording sites located in the hippocampus anterior to −4.5mm and more dorsally than −5mm from bregma were considered as recording from the dorsal hippocampus, electrodes located within the same range dorso-ventrally but posterior to −4.5mm from bregma were considered as located in the intermediate hippocampus, and all electrodes below −5mm dorso- ventrally from bregma were considered to be placed within the ventral hippocampus. No electrodes were placed more anteriorly than −3.8mm from bregma.

To assess the distribution of units recorded along the hippocampal septotemporal axis, the estimated coordinates at which each shank crossed area CA1 were projected onto a 3D centerline model of the hippocampal septotemporal axis, constructed by spline-fitting mid-CA1 coordinates extracted from the same brain atlas and using orthogonal projection onto the local axis tangent.

### Data analysis

Analysis of behavioral and neuronal data was performed using Python3.8, its scientific extension modules -numpy(2.2.6) scipy(1.15.2), matplotlib(3.10.8), and seaborn(0.13.2)- together with custom-written code under Spyder and Jupyter Notebooks.

#### Behavior

##### Pose estimation

The positions of the rats’ body parts in the overhead view (snout, head, back center, and tail base) and facial features in the face camera view (cardinal points of the eye and pupil) were estimated using a U-Net model retrained on manually labeled frames in SLEAP(v1.2.9)^100^.

The same iterative procedure was used for body pose estimation and eye tracking. For each animal, 20 randomly selected frames from videos corresponding to each experimental phase were manually labeled (for eye-tracking, only the solo baseline, shock observation, and recall phases were used). Models were initially trained using the default SLEAP parameters, and predictions were generated on 20 additional randomly selected frames from each video. Mislabelled frames were manually corrected and added to the training set before retraining the model. This process was repeated iteratively until no further manual corrections were required on the sampled frames.

For each body and facial feature, pose estimates were further refined by removing coordinates located outside the arena boundaries or deviating from their positions in the preceding and following frames by more than a predefined threshold (15 cm for body features and 15 pixels for facial features). Missing values occurring in gaps shorter than 500 ms were subsequently filled by linear interpolation.

The resulting features’ coordinates were used to quantify head position, speed, freezing, proximity to the demonstrator’s divider, pupil diameter and eye movement.

##### Freezing characterization

The animal’s position and speed within the arena were estimated using the tracking of the animals’ head defined as the midpoint between the ears. The speed was computed as the absolute value of the gradient vector of position over time, smoothed with a Gaussian kernel of 0.2s bandwidth. Freezing was characterized as time epochs with head speed < 0.1cm/s, merging epochs less than 0.2s apart, and lasting more than 5s. The percentage of time spent freezing in the two contexts during recall subtracted from that spent in the same state during the exploration of the same context alone during baseline, was used to assess the changes in freezing following footshock observation.

The threshold to separate non-recallers from the recallers’ animals was defined as the local minima of a Gaussian kernel density estimation of the difference between the baseline corrected percentage of time freezing in the shock and safe context using Scott’s rules and a bandwidth factor of 0.5.

##### Spatial occupancy

Occupancy maps were computed as 2D histograms of the head positions with 2.5 × 2.5cm bins, normalized by time spent and smoothed with a 2D Gaussian kernel of 5cm bandwidth.

To assess occupancy relative to the demonstrator’s compartment, the occupancy was integrated over the axis perpendicular to the divider, and the time spent in its vicinity was quantified using the percentage of time spent in the 3 bins closest to it.

##### Pupil diameter and eye movement

The pupil size was estimated from the recordings performed with the head-mounted camera as the length of the major axis of an ellipse fitted to the 8 cardinal points on the pupil, and eye movement was estimated as the absolute value of the gradient vector of position over time of the pupil centroid. Both were smoothed with a 40ms Gaussian kernel and z-scored.

Shock observation responses were quantified as the mean values within the 3s window prior to and following the footshock delivery.

#### Electrophysiological recordings

##### Spike sorting

Spike waveforms were automatically clustered into putative single units using Kilosort2.5 (https://github.com/MouseLand/Kilosort) ran from the SpikeInterface framework (0.103.3) (https://github.com/SpikeInterface/spikeinterface), and manually curated using the software package Phy2 (https://github.com/cortex-lab/phy) to identify well-isolated and stable single unit activity clusters.

Were excluded from the analysis units with either of the following:

- an overall average firing rate below 0.1sp/s
- a relative Inter-spike-interval (ISI, 1.5ms) violations rate superior to 0.05
- a fraction of missed spikes, estimated from the waveforms’ amplitude distribution, higher than 5%.

Units with a peak-to-through duration longer than 0.45ms and an averaged autocorrelogram within a 50ms window shorter than 25ms were considered pyramidal units, and the rest of the units were considered to be putative interneurons.

Finally, only interneurons with a mean firing rate of at least 0.5spike/s and pyramidal units with a place field covering more than 125cm² with a field mean rate of at least 0.5spike/s during at least one of the experimental phases were included in the single units analysis.

##### Single units shock observation responses

For every single unit, the firing rate over time was computed with 10ms time bins and smoothed with a 50ms Gaussian kernel. The mean neuron response to shock observation was calculated as the average firing rate during the 1s window following each of the 10 footshock presentations delivered to the demonstrator. The same computation was repeated 1000 times after random circular permutations of the firing rates within a 20s time window centered around each footshock presentation to generate a null distribution. Next, the two-sided Monte Carlo P value was estimated as twice the smallest fraction of the null distribution larger or smaller than the true mean response. The same procedure was repeated around the 10 control shocks delivered to the empty grid.

Units with Monte Carlo P value<0.05 for the footshocks to the demonstrators but not the control shocks (we observed only 4 occurrences of a neuron with P value< 0.05 for both, 3 putative pyramidal neurons from the ventral hippocampus, 1 pyramidal neuron from the intermediate) were classified as responsive to the shock observation. Units were considered to be excited or inhibited by the shock observation if their mean response was respectively larger or smaller than their mean firing rate calculated over the 1s window preceding the footshock presentations.

Finally, the proportions of responsive neurons, separately for excited/inhibited response and putative pyramidal/interneurons within each brain region, were compared to a null distribution of pseudo-random proportions obtained from the concatenated 1000 permutations of firing rates as described above for all neurons of each category.

##### Spatial rate maps

Spatial rate maps were computed separately for all experimental phases, from the position of the animals’ head while moving at a speed of at least 5cm/s and computed as the number of spikes emitted within each 2.5 × 2.5cm position bin over the time spent by the animal within the same bin, and for positions with an occupancy of at least 100ms. The rate maps were further smoothed with a 5cm 2D-Gaussian kernel.

Place fields were defined as continuous bins with a firing rate of at least 20% of the maximum position rate. Field peak rates were estimated as the within-field maximum rate. Spatial information was computed as:

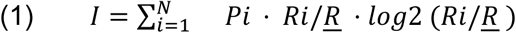

Where:

*N*: total number of spatial bins

*Pi*: occupancy probability of bin *i*

*Ri*: mean firing rate in bin *i*

*<u>R</u>*: overall mean firing rate across all bins

##### Place-shock observation conjunctive coding

To assess whether hippocampal neurons conjunctively encode shock observation and place, we tested the hypothesis that the change in firing rate associated with the shock observation was dependent on the observer’s location relative to each neuron’s place field while footshocks were delivered to the demonstrators.

Shock-observation responses were computed as the difference in mean firing rate computed as described above and normalized by the neuron’s maximum firing rate between the 1s following shock delivery and the 1s preceding it. To account for the effect of the animal’s speed and proximity to the demonstrator’s on the firing rate, the corrected shock-observation-response was obtained by regressing out the mean speed and distance to the demonstrator’s compartment during the 1s of footshock delivery.

To estimate baseline spatial rate during the 1s of footshock delivery, we estimated the time spent by the animal within each spatial bin used to compute spatial rate maps. For each neuron, we used these values to estimate the mean spatial rate from the spatial rate maps, normalized by the maximum spatial rate, computed from the exploration of the observer’s compartment while alone and in the shock context during baseline.

Finally, as an additional control for the variation in firing rate specifically associated with times of footshock delivery, we estimated for each neuron the mean rate difference as described for shock- observation epochs, but for pseudo-randomly selected time epochs -outside of footshock delivery- and while matching the animal location during each shock-observation event.

##### Shock observation moments discrimination analysis

For each animal independently, we trained and tested a linear classifier to discriminate between shock-observation and matched-control moments using speed- and proximity-corrected responses from single neurons, as described above for the Place–shock observation conjunctive coding analysis. Discrimination performance was assessed using a leave-one-trial-out cross-validation procedure performed recursively across trials, yielding an area-under-the-curve (AUC) score quantifying discriminability between shock and control moments.

This analysis was conducted for neurons recorded across the entire hippocampus as well as for neurons restricted to individual hippocampal subregions. To equilibrate neuron counts across animals and regions, we randomly subsampled neurons 100 times using a fixed number of neurons per animal and region, defined by the smallest neuron count across conditions, with a minimum of five neurons per sample set.

##### Changes in spatial representations

To characterize changes in spatial representations, for each neuron, we computed the correlation distance between pairs of spatial rate maps as:

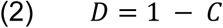

Where:

*D*: distance

*C*: Pearson correlation coefficient between the two spatial maps.

To estimate the effect of shock observation on the spatial representations, we computed the distance between the rate maps estimated from the solo exploration of the observer’s compartment during the recall and baseline, separately for the shock and safe context.

To estimate changes in context discrimination, distances between the spatial rate maps in the safe and shock context were computed separately for the solo exploration of the observer’s compartment in the baseline and recall phase.

Comparisons for which rate maps overlapped with less than half the total surface area of the observer’s compartment were excluded.

##### Sharp Wave Ripples detection

For the reactivation analyses, sharp-wave ripples (SWRs) were detected from local field potentials recorded in CA1 within each of the dorsal, intermediate and ventral hippocampal subregions. We refer to these detection sites as dCA1, iCA1 and vCA1; this notation indicates the anatomical location of the LFP channels used for SWR detection, not a restriction of all simultaneously recorded single units to CA1. CA1 channels were used because sharp waves reflect excitatory CA3 input to CA1 through Schaffer collaterals, and the associated ripple-frequency oscillation is most clearly detected in CA1 local field potentials101,102. The local field potentials recorded from 1-5 electrodes recorded in CA1 within the dorsal, intermediate and ventral hippocampus were hand-picked after visual inspection for the presence of large SWRs, separately in the right and left hemispheres. The signals were first phase-shift corrected, then re-referenced using a common median reference from all recorded channels on a single probe and downsampled to 1KHz. The local field potentials were then filtered within the ripple frequency band (150-250Hz). Because ripple peak frequency has been reported to decrease along the dorsoventral axis^101,102^ this fixed band may have led to comparatively conservative SWR detection in the ventral hippocampus. A ripple envelope was computed as the absolute value of the Hilbert transform of the ripple-filtered signal, averaged across the recording sites and smoothed with a Gaussian kernel (bandwidth 10 ms). The ripple envelope was detrended using a moving median filter with 3s window. Ripples starts and ends were defined as the times when the envelope exceeded a low threshold of *μ* + 1*σ* while exceeding a high threshold of *μ* + 5*σ* in between, where *μ* and *σ* represent respectively the mean and standard deviation of the detrended envelope. Ripples events with a duration of less than 40ms and more than 250ms were excluded.

##### Post-experience neuronal reactivations

For each animal, the explained variance was computed as the partial correlation between the cell-pair coherence between cell pairwise correlation matrix during exploration and the post-learning rest phase, given the known cell-pair coherence between exploration and post-learning rest with pre-learning rest.:

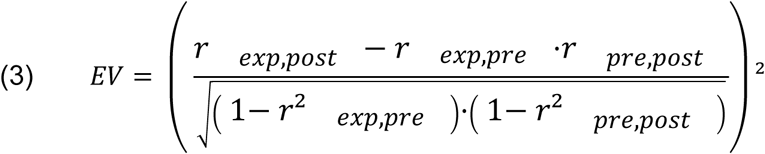

Where:

*EV*: explained variance

*r _x_*_,*y*_: cell-pair coherence between phase x and y

The reversed explained variance was computed as the partial correlation between cell-pair coherence between exploration and pre-learning rest, given the known coherence between experience and pre-learning rest with post-learning rest by swapping pre-exploration and post-experience rest phases in the equation above.

Both explained variance were computed separately for the entire baseline exploration of the observer’s compartment, alone and with the demonstrator present in the safe and shock context, as well as the shock observation phase separately.

Pairwise correlation matrices for all phases were computed from the z-scored firing rates, computed in 100ms time bins and smoothed with a 0.2s gaussian kernel. For all exploration phases, time bins overlapping with a SWR detection in any of the hippocampal sub-regions from any hemisphere were excluded before computing Pearson correlations. The opposite logic was followed for all the rest phases, for which pairwise Pearson correlations were computed only from time bins overlapping with a SWR detection.

##### Inter-subregion SWR cross-correlation

For each animal, cross-correlograms with a 20ms bin size were computed between SWR peak amplitude times for events detected during the post-learning rest period. For this analysis and when applicable, detections from both hemispheres were merged.

#### Statistical analysis

Frequentist and Bayesian statistical testing was performed using Python 3.8 together with the scientific extension modules pandas(2.3.3), pingouin(0.6.1), scipy(1.15.2) and statsmodels(0.14.6). In the case of the linear mixed model analysis, Bayesian modeling was performed in R 4.5 using the brms package^103^ and Bayes factors were computed using the bayestestR package^104^.

Differences in freezing time, time near the demonstrator, pupil diameter, eye movements, neural firing properties, shock-moment discrimination AUC, and SWR explained variance were assessed using paired two-sided t-tests for within-animal comparisons (safe vs. shock context; pre- vs. post-shock) and two-sided independent-samples t-tests for between-group comparisons (recaller vs. non-recaller; excited vs. non-excited units). Where multiple related comparisons were performed within the same analysis family, P values were corrected using the Holm method; corrected P values are reported as such in the corresponding figure legends and text. Normality was assessed using D’Agostino and Pearson’s omnibus test. This assumption was met for the main effects (discrimination AUC and SWR explained-variance comparisons). For contextual freezing, as reported, the data deviated from normality; a two-sided Wilcoxon signed-rank test nonetheless confirmed the same pattern of statistical significance as the paired t-test, and the Student’s t-test is reported to allow direct comparison with the corresponding Bayesian estimate. For linear mixed models, normality of model residuals was assessed by visual inspection of quantile–quantile (Q-Q) plots.

Differences in the proportion of units classified as excited, inhibited, or unresponsive to shock observation across the dorsal, intermediate, and ventral hippocampus were assessed using a χ² test of proportions.

The effect of distance from the demonstrator’s compartment on rat occupancy and single-cell mean normalized firing rate was tested using an analysis of covariance (ANCOVA) with the following model:

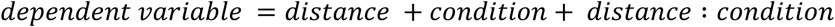

Where:

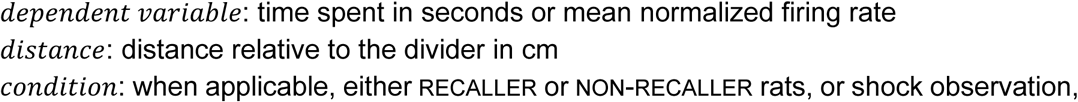

excited or not single units.

To test the main and interaction fixed effects of individual neurons’ spatial firing rate at the time of footshock delivery on single-neuron changes in spiking rate, we used the following linear mixed model:

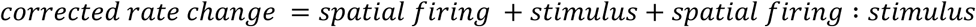

Where:

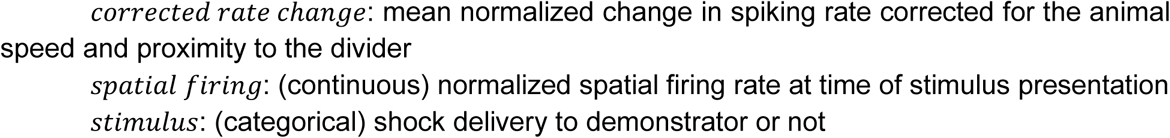

We tested for the effect of rats’ recall status (RECALLER and NON-RECALLER) and single neurons’ response to shock observation (excited or not) separately by adding these variables

To test the main and interaction effects of context or experimental phase and the animal’s recall status (recaller vs. non-recaller) on the correlation distance between spatial rate maps of single units, we used the following linear mixed model:

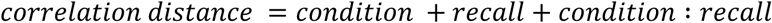

Where:

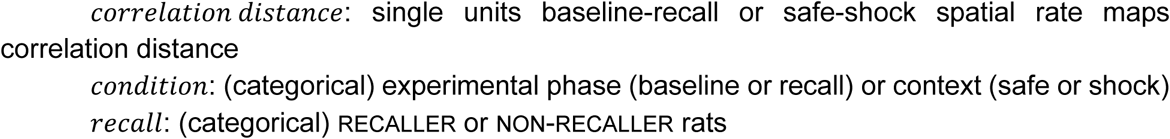

For all linear mixed models, rat identity was included as a random intercept to account for inter-individual variability, and fixed-effect coefficients were evaluated using two-sided one-sample t-tests against zero. For the Bayesian estimations, we used Cauchy priors centered on 0 and with a scale of 0.2.

For all Bayesian inferences, we consider BF10s of 1 as indicating indifference for the support of either the null or alternative hypothesis; BF10s < 1 as suggesting evidence in favor of the null hypothesis, and BF10s > 1 as indicating evidence in favor of the alternative hypothesis^105,106^. Following traditionally used thresholds, we consider BF10<1/3 or BF10>3 as moderate evidence for the absence or presence of an effect, BF10<1/10 or BF10>10 as strong evidence, and 1/3<BF10<3 as circumstantial evidence.

## Data availability

Source data are provided with this paper and via Zenodo (DOI: 10.5281/zenodo.22677727). Raw electrophysiological and behavioral data generated in this study will be made available without restrictions upon request to the first corresponding author (FML).

## Code availability

All analyses were conducted using custom code in Python3.8 and R. Custom analysis code is publicly available in a dedicated repository available in GitHub (https://github.com/FredMichon/HippocampalShockObservationResponses) and Zenodo (DOI: 10.5281/zenodo.22677727).

## Acknowledgements

We would like to thank Francesco Battaglia and Baptiste Mahéo for their feedback on the analysis of the reactivation of experience-related patterns of spiking activity. We thank Lina Asperl for her help with the pose estimation of the eye and pupil diameter. Finally, we are thankful to Moritz Koch and Ákos Babiczky for their support with histology.

## Authors contribution

Conceptualization: FML, VG, CK

Data curation: FML

Formal analysis: FML, CK

Funding acquisition: FML, VG, CK

Investigation: FML

Project administration: FML

Resources: FML, CK

Visualization: FML

Writing – original draft: FML

Writing – review and editing: FML, VG, CK

## Supplementary Figures and Tables

**Supplementary Table 1:**
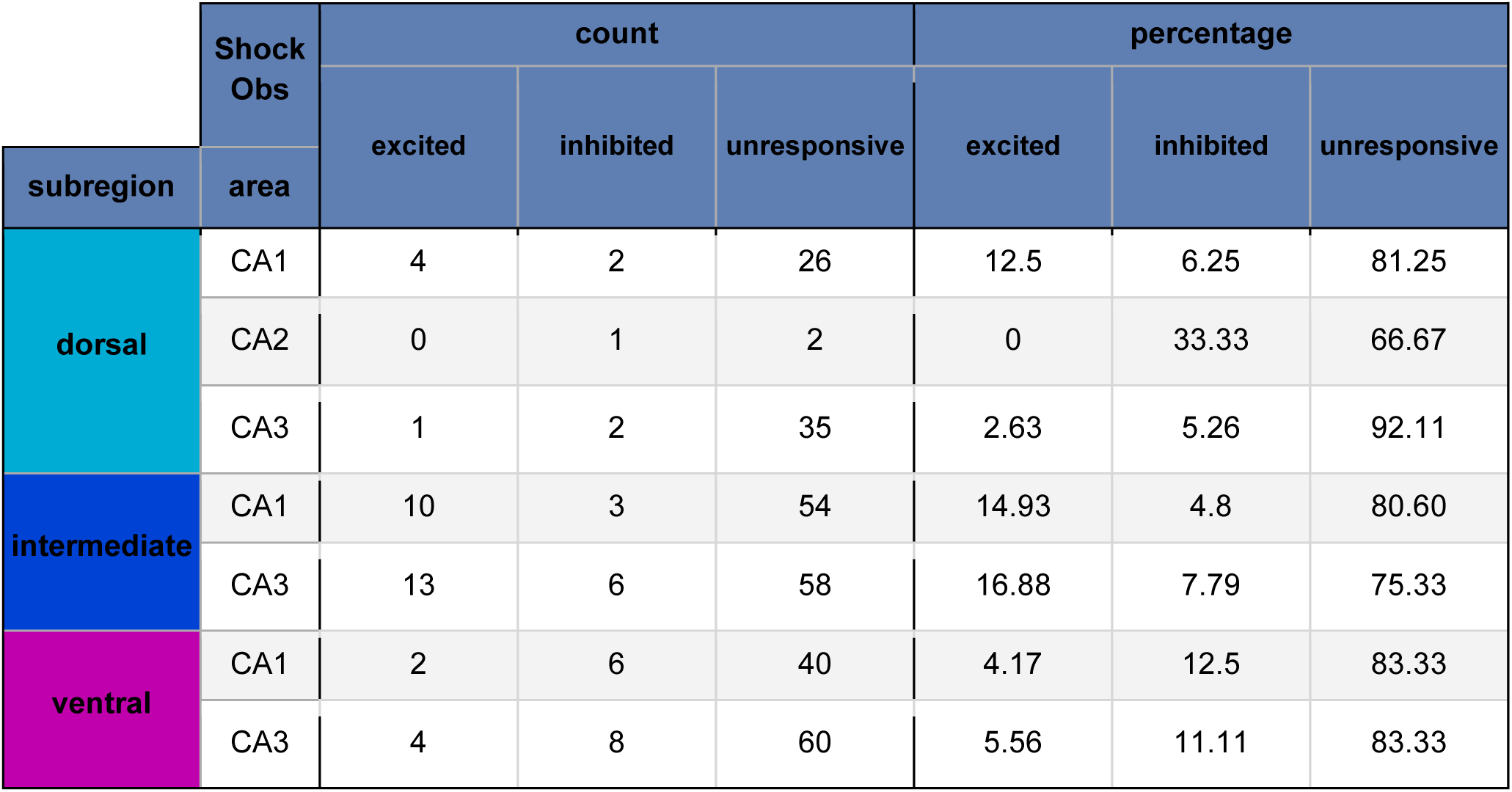
Quantification of pyramidal neurons by estimated recording location and response at the moment of shock observation.

**STable 2:**
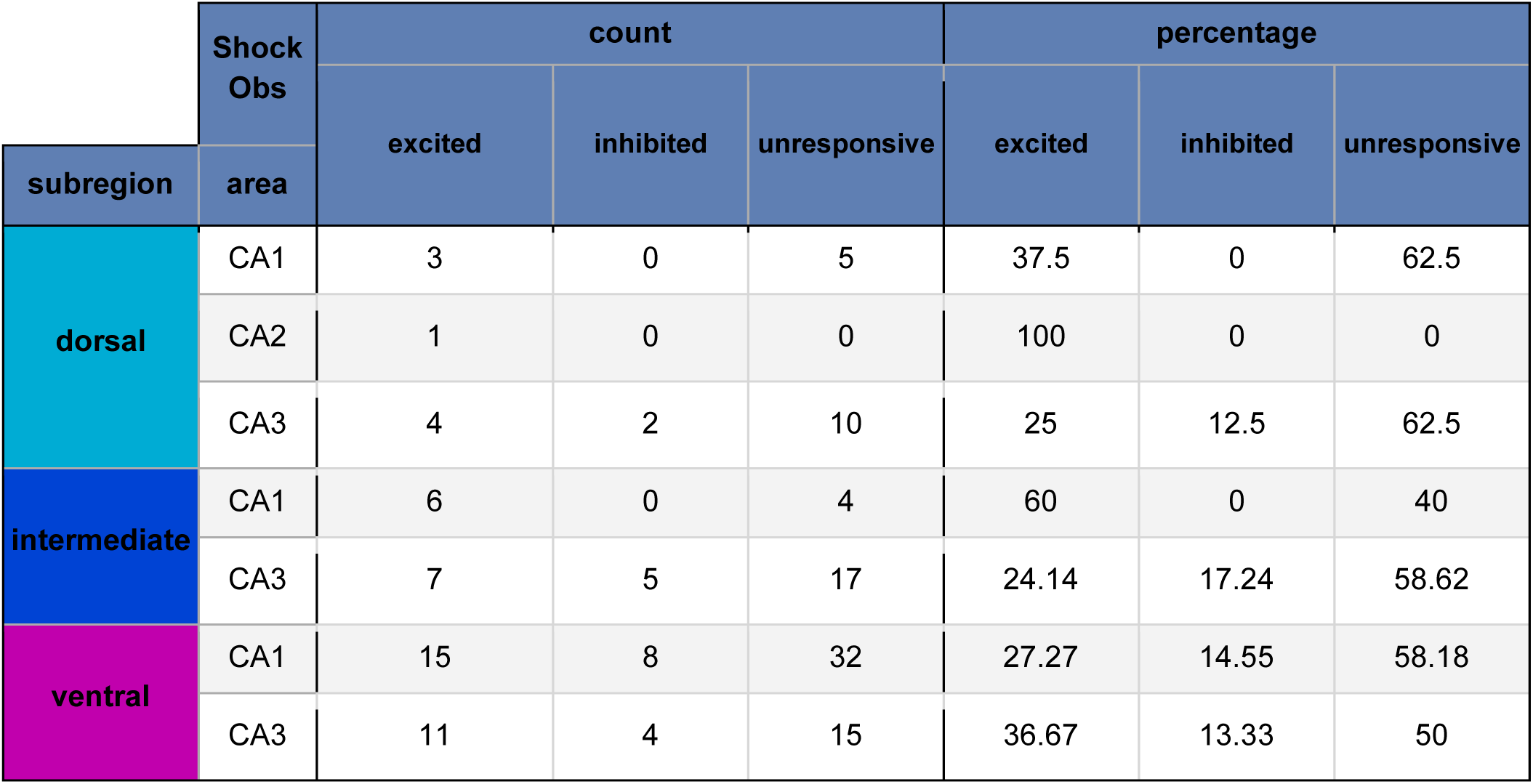
Quantification of interneurons by estimated recording location and response at the moment of shock observation.

**Supplementary Figure 1:**
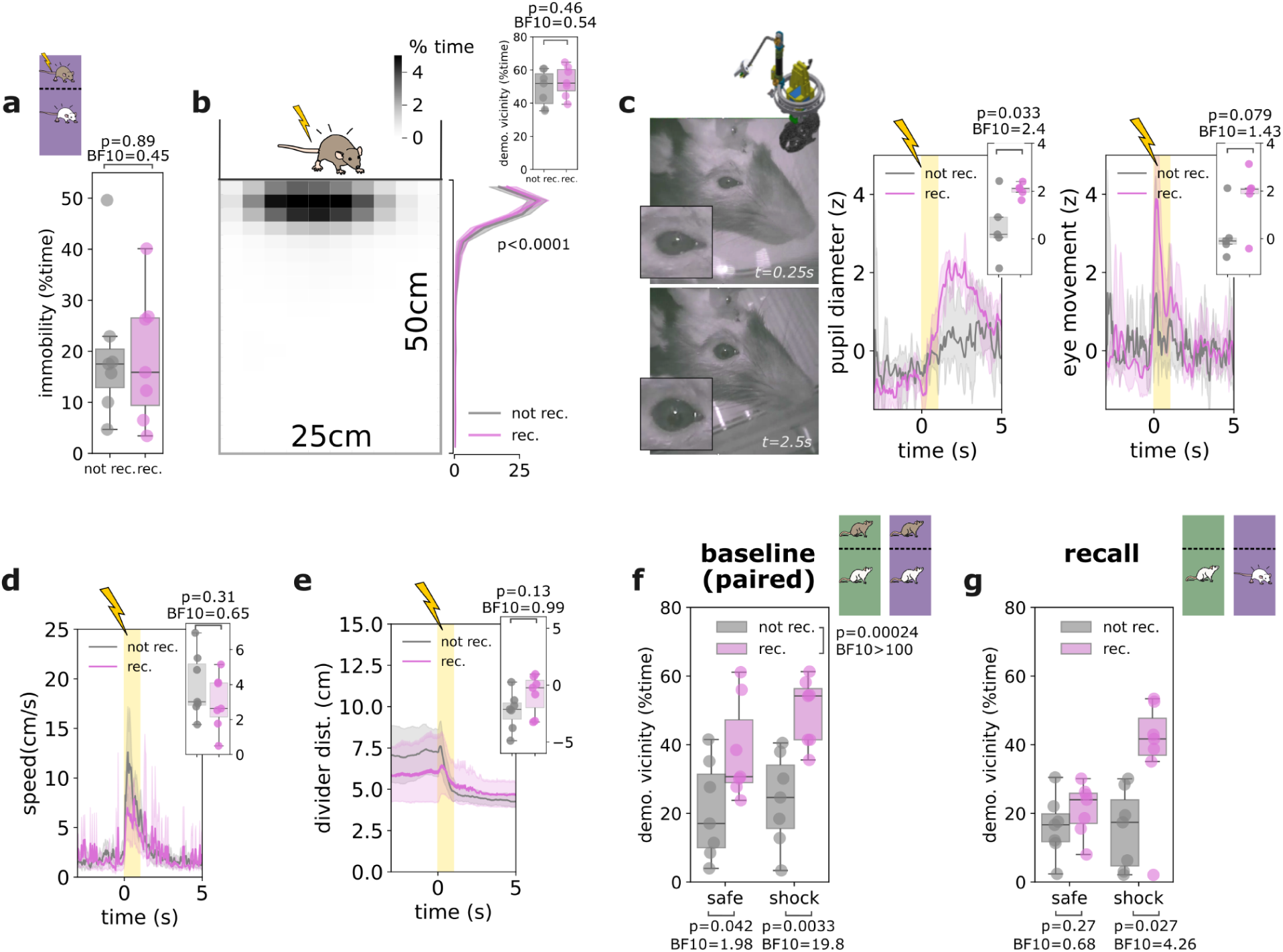
Behavioral correlates of contextual vicarious fear learning. **a.** No observed difference in time spent freezing during the shock observation between RECALLER (rec., in pink) or NON-RECALLER (not rec., in grey) rats. **b.** During the shock observation, all animals spent a majority of time with their head at the vicinity of the divider separating the demonstrator’s compartment. Left panel, averaged occupancy map for all rats. Right panel, average occupancy relative to the distance to the divider separately for RECALLER (rec.) or NON-RECALLER (not rec.) observer rats. Inlet, average percentage of time spent with the head within 7.5cm of the demonstrator’s divider separately for RECALLER (rec.) or NON-RECALLER (not rec.) observer rats. **c.** Stronger pupil and eye-ball responses for RECALLER (rec.) or NON-RECALLER (not rec.) rats. Left panel shows two example frames recorded from the face of a single observer 0.25s prior (top) and 2.5s following footshock delivery to the demonstrator. Right, averaged z-scored pupil diameter and eye movement around footshock delivery to the demonstrator, separately for RECALLER (rec.) or NON-RECALLER (not rec.) observer rats. Inlet boxplots represent the mean difference between the 3s following and preceding the onset of shock delivery. **d-e**. Similar orienting response for all animals during shock observation. Averaged speed (**d**) and distance to the demonstrator’s divider (**e**) around footshock delivery to the demonstrator, separately for RECALLER (rec.) or NON-RECALLER (not rec.) observer rats. Inlet boxplots represent the mean difference between the 3s following and preceding the onset of shock delivery. **f-g.** Higher propensity to interact with the demonstrator for rats that later recalled the shock-context association. Percentage of time spent at the vicinity of the demonstrator’s compartment in the safe and shock contexts, separately for RECALLER (rec.) or NON-RECALLER (not rec.) observer rats, during baseline with the demonstrator **(f)** and during recall **(g)**. Boxplots center line represents the median, the box limits the 25th and 75th percentiles, and the whiskers extend to values up to 1.5 x the interquartile range; points indicate values for individual animals. Plain lines represent averages, and corresponding shaded areas the 95% confidence interval estimated from 500 bootstrapping iterations. Yellow shaded areas mark the shock- delivery duration. Student T-test P value and Bayesian BF10 are reported, except for the right panel in b, for which the P value reported is that of the effect of the distance to the demonstrator’s divider of an ANCOVAdistanceXlearning. For all panels data displayed are from N = 14 rats (7 RECALLERS, 7 NON-RECALLERS), except in (c) (N = 10 rats (5 RECALLERS, 5 NON-RECALLERS). Source data are provided as a Source Data file.

**Supplementary Figure 2:**
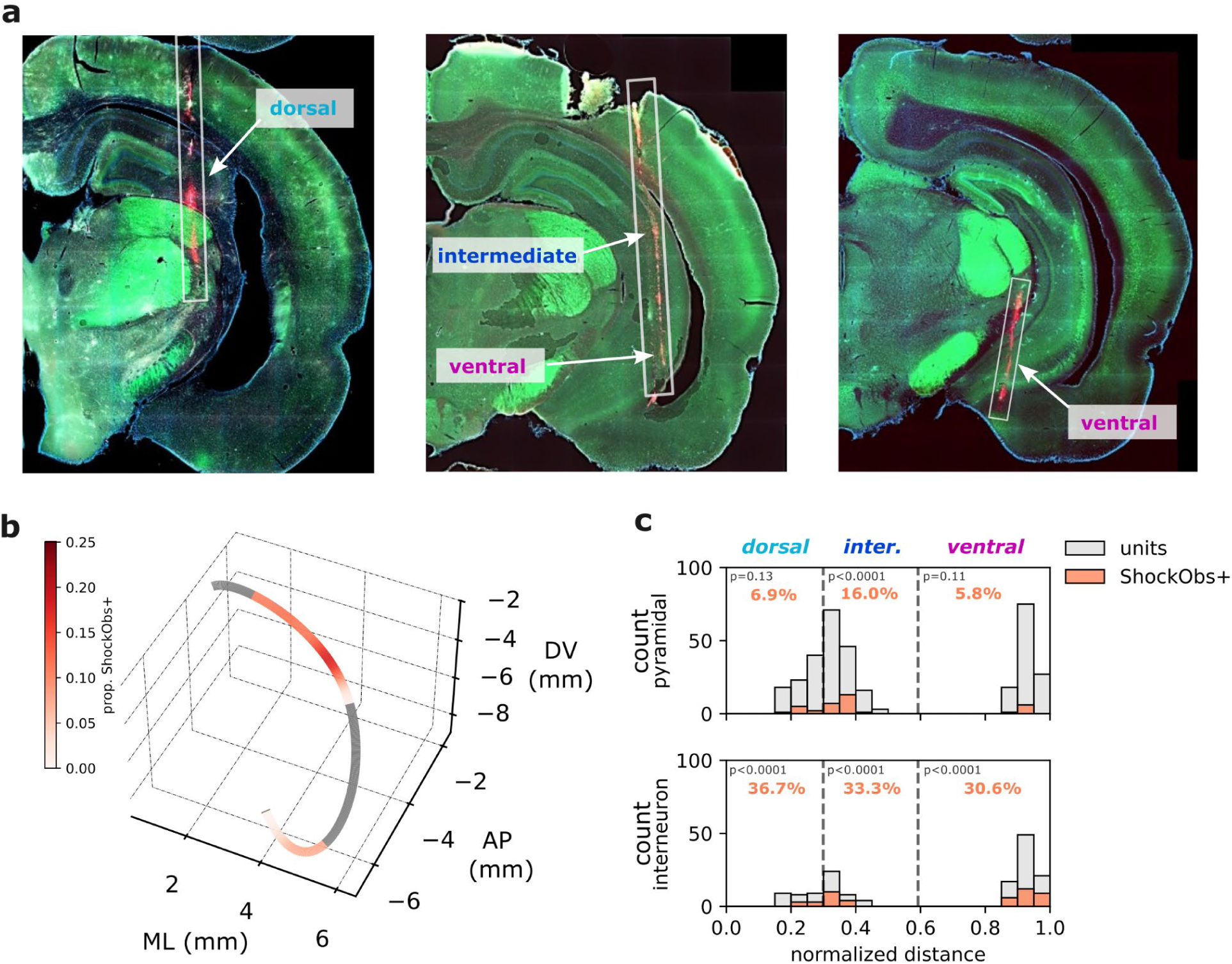
Anatomical mapping of hippocampal recording sites. **a.** Example coronal sections used to reconstruct implantation sites, with probe shank tracks crossing the hippocampal principal sheets at coordinates classified as corresponding to the dorsal, intermediate or ventral hippocampus. Images were processed in QuPath (v0.2.3), where each color channel was thresholded individually. **b.** Proportion of shock observation responsive pyramidal neurons mapped onto a 3-D reconstruction of the hippocampal septo-temporal axis along mid-area CA1. Gray segments indicate portions of the septotemporal axis that were not sampled. **c.** Distribution of all (units, in grey) pyramidal (top panel, N = 337 neurons) or interneurons (bottom panel, N = 149) and shock observation responsive units ( shockobs+, orange) along the septotemporal axis as shown in B. Dashed lines mark the border used to define dorsal, intermediate and ventral hippocampal subregions. P values are two-sided Monte Carlo P values for testing whether the number of responsive neurons exceeds that expected by chance estimated from random temporal shifting. Source data are provided as a Source Data file.

**Supplementary Figure 3:**
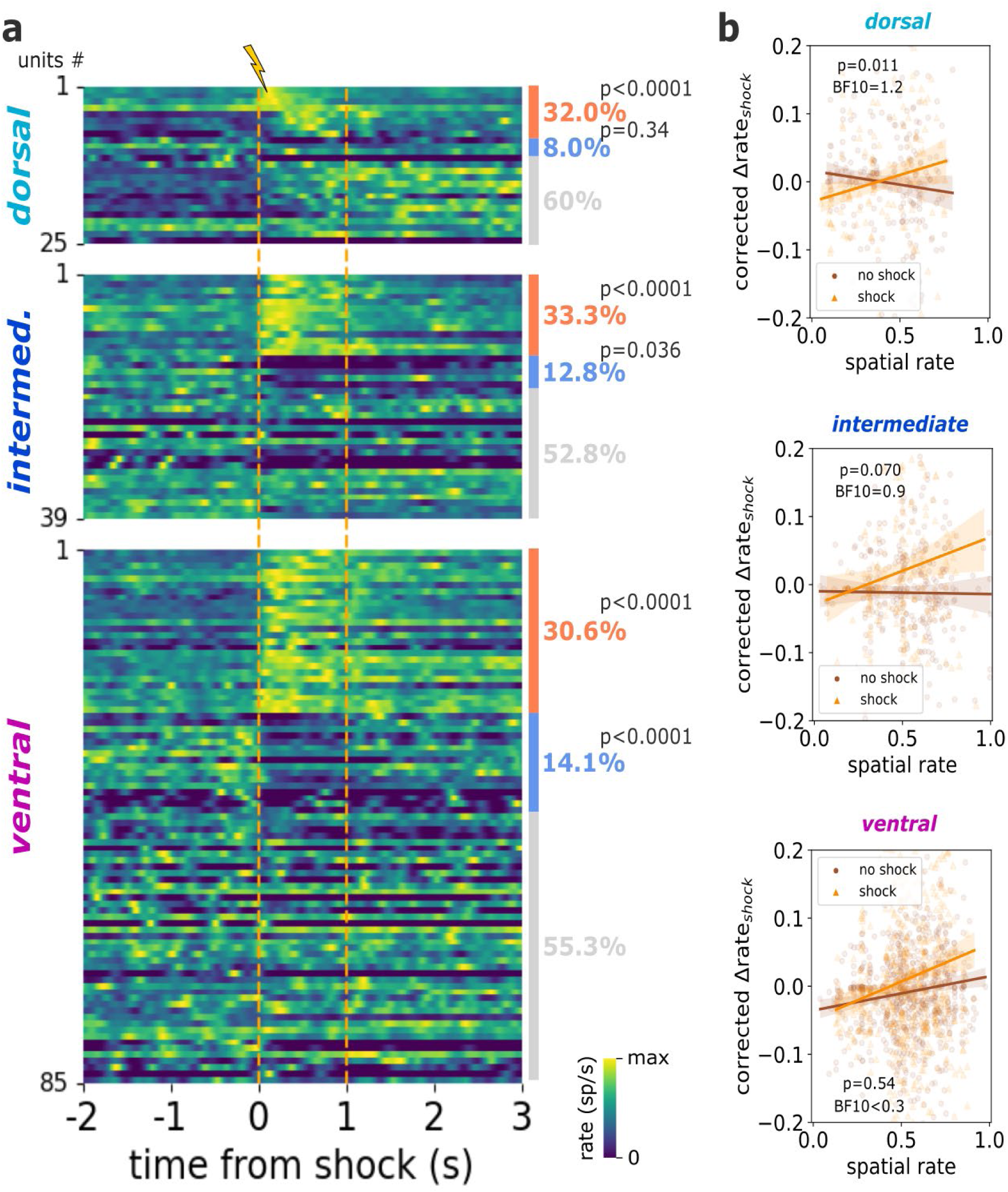
Interneurons response to shock observation. **a.** Heatmap representing the mean firing rate of single putative interneuron split by hippocampal sub-region (dorsal, intermediate, ventral). For each sub-region, neurons are ordered from top to bottom as follows: significantly excited (orange), significantly inhibited (blue) and finally not significantly modulated (grey) single neurons. Overall percentages are indicated on the right side of the heatmaps. Orange dashed vertical lines represent the onset and offset of footshock delivery. **b.** For each hippocampal subregion, linear regression between corrected neuronal response for speed and distance to the demonstrator’s compartment and estimated basal spatial firing rate at times of footshock delivery (gold) or matched control-moments without shocks (sienna). In (**a**) Monte Carlo P value for testing whether the number of responsive neurons exceeds that expected by chance estimated from random temporal shifting. In (**b)**, mean regression line with 95% confidence interval. Dots represent single trial responses. P value and BF10 for the linear mixed model’s fixed effect of the interaction: baseline spatial rate X footshock delivery times. Source data are provided as a Source Data file.

**Supplementary Figure 4:**
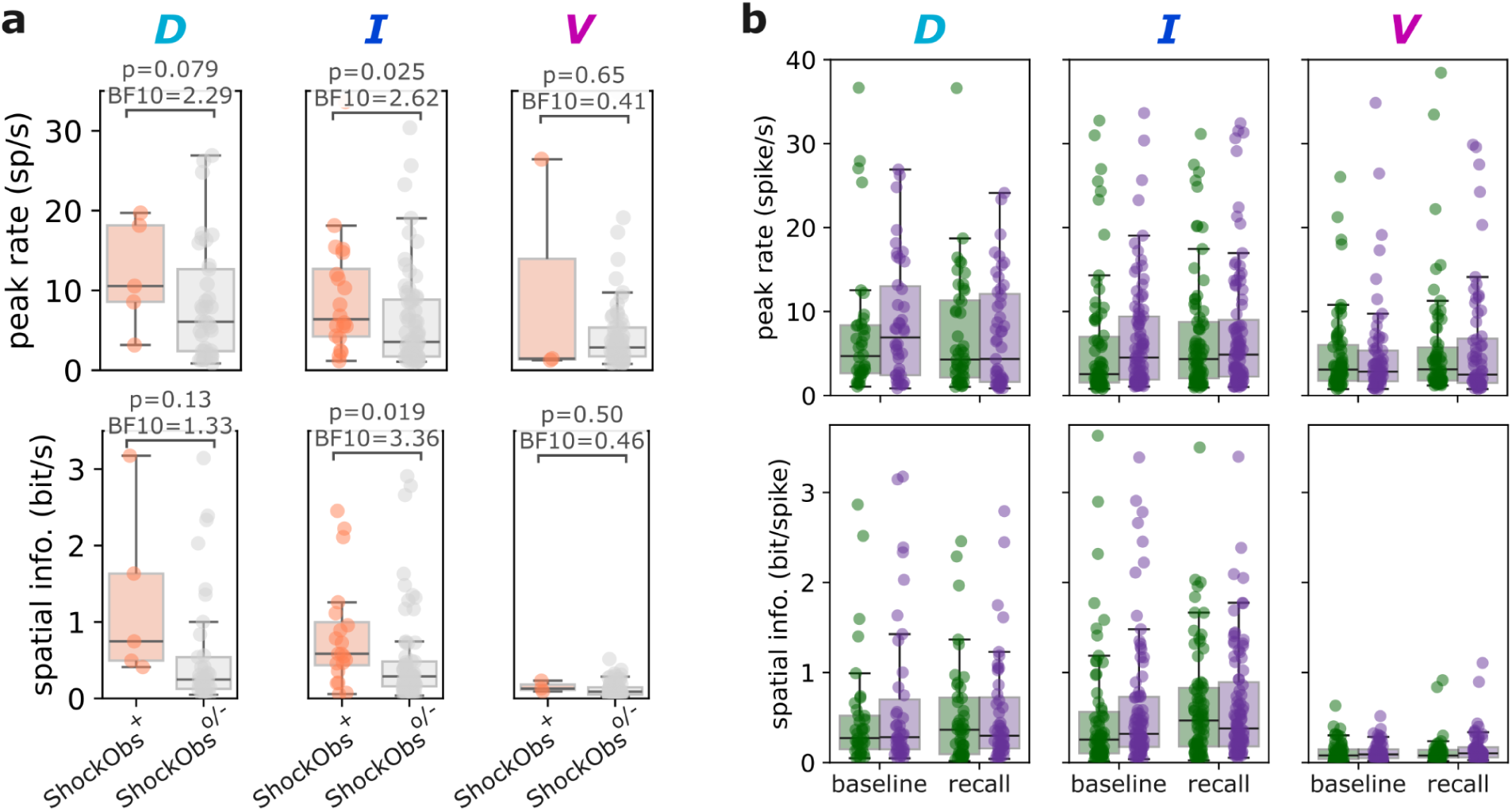
Spatial firing properties. **a.** Spatial tuning properties of putative pyramidal neurons during solo baseline split by hippocampal subregion (dorsal, D, intermediate, I, and ventral, V) separately for neurons significantly excited (ShockObs^+^, in orange, N = 5/23/6 neurons in D/I/V) or not (unresponsive and inhibited, ShockObs^o/-^, in grey, N = 68/121/114 neurons in D/I/V) by the shock observation. Top row, maximum field rate. Bottom row, spatial information. **b.** For all putative pyramidal neurons, peak field rate (top) and spatial information (bottom) for spatial rate maps estimated from the exploration of the observer’s compartment in the safe (green) and shock (purple) context, and during solo baseline and recall. Boxplots center line represents the median, the box limits the 25th and 75th percentiles, and the whiskers extend to values up to 1.5 x the interquartile range; points indicate values for individual neurons. In **a**, uncorrected P values and BF10 from two-sided student T-test. In **b,** Top, two-way ANOVA (context × phase), in dorsal: context (F(1,258)=0.74, p=0.39, η²p=0.003), phase (F(1,258)=0.17, p=0.68, η²p<0.001), interaction (F(1,258)=0.02, p=0.88, η²p<0.001); in intermediate: context (F(1,543)=3.67, p=0.056, η²p=0.007), phase (F(1,543)=2.96, p=0.086, η²p=0.005), interaction (F(1,543)=0.02, p=0.88, η²p<0.001); context (F(1,417)=0.08, p=0.77, η²p<0.001), phase (F(1,417)=0.19, p=0.67, η²p<0.001), or interaction (F(1,417)=0.89, p=0.35, η²p=0.002). Bottom, two-way ANOVA (context × phase), in dorsal: context (F(1,276)=0.67, p=0.41, η²p=0.002), phase (F(1,276)=0.88, p=0.35, η²p=0.003), interaction (F(1,276)=0.29, p=0.59, η²p=0.001); in intermediate: context (F(1,572)=0.31, p=0.58, η²p<0.001), phase (F(1,572)=2.44, p=0.12, η²p=0.004), interaction (F(1,572)=0.32, p=0.57, η²p<0.001); in ventral: context (F(1,476)=1.05, p=0.31, η²p=0.002), phase (F(1,476)=0.58, p=0.45, η²p=0.001), or interaction (F(1,476)=0.81, p=0.37, η²p=0.002). Source data are provided as a Source Data file.

**Supplementary Figure 5:**
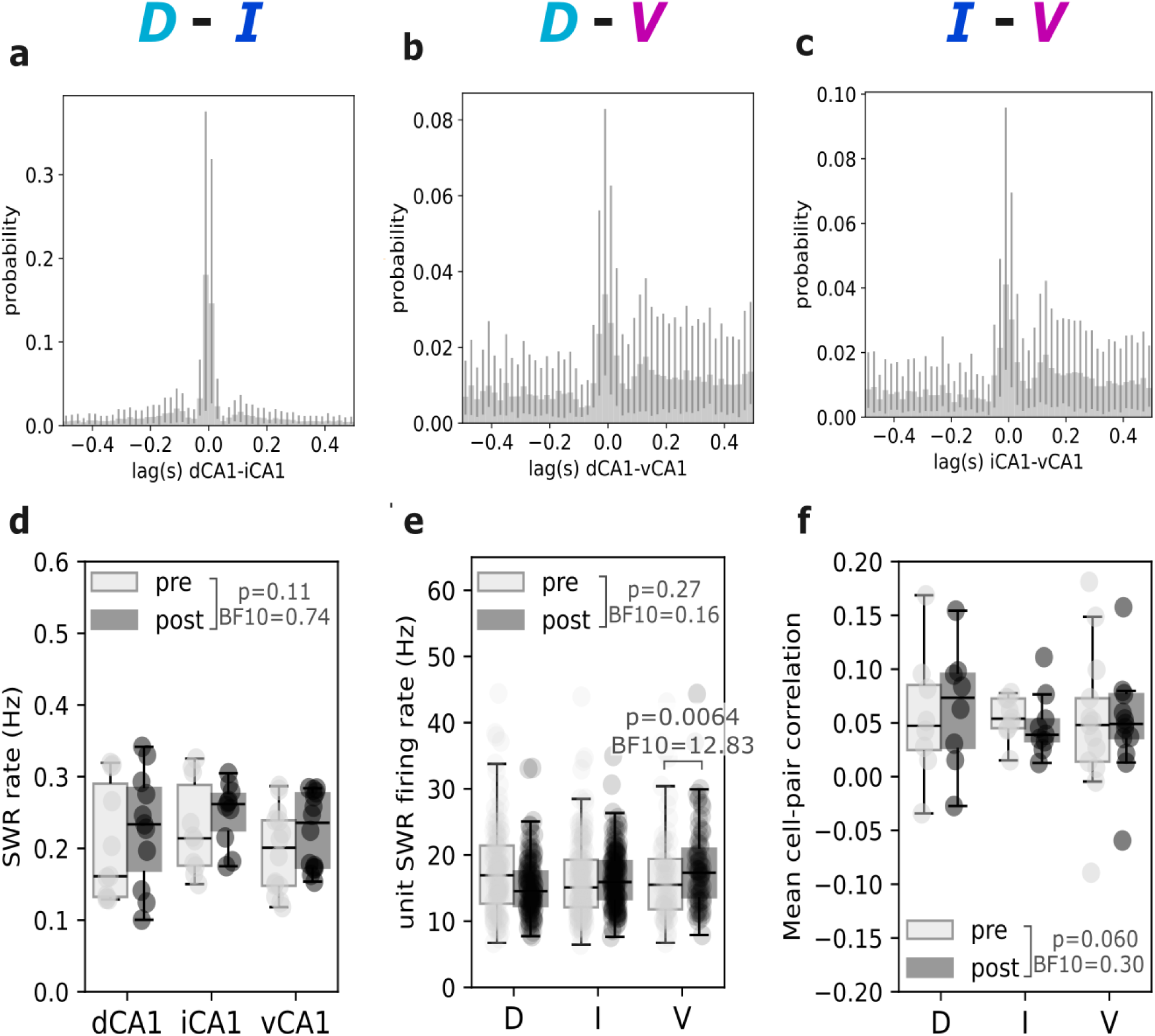
SWR-related activity across hippocampal subregions. Stronger dorsal-intermediate SWR synchrony compared to dorsal-ventral and intermediate- ventral SWR occurrences during the post-rest phase. Mean cross-correlograms for the occurrence of SWRs detected in the dorsal and intermediate hippocampus (N = 7 rats) (**a**), dorsal and ventral hippocampus (N = 11 rats) (**b**), and intermediate and ventral hippocampus (N = 10 rats) (**c**). **d.** Mean rates of sharp-wave ripples occurrence detected in the dorsal (dCA1, N = 11 rats), intermediate (iCA1, N=10) and ventral (vCA1, N = 14 rats) hippocampus, computed for each rat during the pre-shock (light grey) and post-shock (dark grey) rest phases. **e.** Mean single-unit firing rate of recorded pyramidal neurons during sharp-wave ripple occurrences detected in the pre-shock and post-shock rest phases, and separated for units recorded from the dorsal (D, N= 73 neurons), intermediate (I, N = 144 neurons) and ventral (V, N = 120 neurons) hippocampus. **f.** For each rat, the mean pyramidal cell pair firing rate correlation during sharp-wave ripples detected in the pre-shock and post-shock rest phases, and separated for units recorded from the dorsal (D, N = 8 rats), intermediate (I, N = 9 rats) and ventral (V, N = 13 rats) hippocampus. Error bars represent the 95% confidence interval estimated from 500 bootstrapping iterations. In **d-f**, Boxplots center line represents the median, the box limits the 25th and 75th percentiles, and the whiskers extend to values up to 1.5 x the interquartile range; points indicate values for individual animals. In **d**, two-way ANOVA (subregion × sleep phase): subregion (F(2,64)=1.13, p=0.33, η²p=0.034), phase (F(1,64)=2.51, p=0.12, η²p=0.038), interaction (F(2,64)=0.02, p=0.98, η²p<0.001). In **e**, two-way ANOVA (subregion × sleep phase): subregion (F(2,660)=2.27, p=0.10, η²p=0.007), phase (F(1,660)=1.25, p=0.26, η²p=0.002), interaction (F(2,660)=5.12, p=0.006, η²p=0.015). In **f**, two-way ANOVA (subregion × sleep phase): phase (F(1,54)=0.27, p=0.61, η²p=0.005), subregion (F(2,54)=0.11, p=0.89, η²p=0.004), interaction (F(2,54)=0.23, p=0.80, η²p=0.008). In **d-f**, corrected P value and Bayesian BF10 from st hoc ANOVA student T-test are reported. Source data are provided as a Source Data file.

## References

1. Ferretti, V. & Papaleo, F. Understanding others: Emotion recognition in humans and other animals. Genes Brain Behav. 18, e12544 (2019).

2. Atsak, P. et al. Experience modulates vicarious freezing in rats: a model for empathy. PloS One 6, e21855 (2011).

3. Jeon, D. et al. Observational fear learning involves affective pain system and Cav1.2 Ca2+ channels in ACC. Nat. Neurosci. 13, 482–488 (2010).

4. Keysers, C., Knapska, E., Moita, M. A. & Gazzola, V. Emotional contagion and prosocial behavior in rodents. Trends Cogn. Sci. 26, 688–706 (2022).

5. Terranova, J. I., Yokose, J., Osanai, H., Ogawa, S. K. & Kitamura, T. Systems consolidation induces multiple memory engrams for a flexible recall strategy in observational fear memory in male mice. Nat. Commun. 14, 3976 (2023).

6. Carrillo, M. et al. Emotional Mirror Neurons in the Rat’s Anterior Cingulate Cortex. Curr. Biol. 29, 1301–1312.e6 (2019).

7. Allsop, S. A. et al. Corticoamygdala Transfer of Socially Derived Information Gates Observational Learning. Cell 173, 1329–1342.e18 (2018).

8. Kim, S.-W. et al. Hemispherically lateralized rhythmic oscillations in the cingulate-amygdala circuit drive affective empathy in mice. Neuron 111, 418–429.e4 (2023).

9. O’Keefe, J. & Nadel, L. The Hippocampus as a Cognitive Map. (Clarendon Press, Oxford, 1978).

10. Huxter, J., Burgess, N. & O’Keefe, J. Independent rate and temporal coding in hippocampal pyramidal cells. Nature 425, 828–832 (2003).

11. Lu, L. et al. Impaired hippocampal rate coding after lesions of the lateral entorhinal cortex. Nat. Neurosci. 16, 1085–1093 (2013).

12. Takahashi, S. Episodic-like memory trace in awake replay of hippocampal place cell activity sequences. eLife 4, e08105 (2015).

13. Sanders, H. et al. Temporal coding and rate remapping: Representation of nonspatial information in the hippocampus. Hippocampus 29, 111–127 (2019).

14. Terada, S., Sakurai, Y., Nakahara, H. & Fujisawa, S. Temporal and Rate Coding for Discrete Event Sequences in the Hippocampus. Neuron 94, 1248–1262.e4 (2017).

15. Ferbinteanu, J., Shirvalkar, P. & Shapiro, M. L. Memory Modulates Journey- Dependent Coding in the Rat Hippocampus. J. Neurosci. 31, 9135–9146 (2011).

16. Latuske, P., Kornienko, O., Kohler, L. & Allen, K. Hippocampal Remapping and Its Entorhinal Origin. Front. Behav. Neurosci. 11, (2018).

17. Moser, M.-B., Rowland, D. C. & Moser, E. I. Place Cells, Grid Cells, and Memory. Cold Spring Harb. Perspect. Biol. 7, a021808 (2015).

18. Allen, K., Rawlins, J. N. P., Bannerman, D. M. & Csicsvari, J. Hippocampal Place Cells Can Encode Multiple Trial-Dependent Features through Rate Remapping. J. Neurosci. 32, 14752–14766 (2012).

19. Alme, C. B., et al. Place cells in the hippocampus: Eleven maps for eleven rooms. Proc. Natl. Acad. Sci. 111, 18428–18435 (2014).

20. Wang, M. E. et al. Long-Term Stabilization of Place Cell Remapping Produced by a Fearful Experience. J. Neurosci. 32, 15802–15814 (2012).

21. Kentros, C. G., Agnihotri, N. T., Streater, S., Hawkins, R. D. & Kandel, E. R. Increased Attention to Spatial Context Increases Both Place Field Stability and Spatial Memory. Neuron 42, 283–295 (2004).

22. Matsuo, N. Irreplaceability of Neuronal Ensembles after Memory Allocation. Cell Rep. 11, 351–357 (2015).

23. Tanaka, K. Z. et al. Cortical Representations Are Reinstated by the Hippocampus during Memory Retrieval. Neuron 84, 347–354 (2014).

24. Robinson, N. T. M. et al. Targeted Activation of Hippocampal Place Cells Drives Memory-Guided Spatial Behavior. Cell 183, 1586–1599.e10 (2020).

25. Davidson, T. J., Kloosterman, F. & Wilson, M. A. Hippocampal Replay of Extended Experience. Neuron 63, 497–507 (2009).

26. Ego-Stengel, V. & Wilson, M. A. Disruption of ripple-associated hippocampal activity during rest impairs spatial learning in the rat. Hippocampus 20, 1–10 (2010).

27. Foster, D. J. & Wilson, M. A. Reverse replay of behavioural sequences in hippocampal place cells during the awake state. Nature 440, 680–683 (2006).

28. Girardeau, G., Benchenane, K., Wiener, S. I., Buzsáki, G. & Zugaro, M. B. Selective suppression of hippocampal ripples impairs spatial memory. Nat. Neurosci. 12, 1222–1223 (2009).

29. Gridchyn, I., Schoenenberger, P., O’Neill, J. & Csicsvari, J. Assembly-Specific Disruption of Hippocampal Replay Leads to Selective Memory Deficit. Neuron 106, 291–300.e6 (2020).

30. Lee, A. K. & Wilson, M. A. Memory of Sequential Experience in the Hippocampus during Slow Wave Sleep. Neuron 36, 1183–1194 (2002).

31. Wilson, M. A. & McNaughton, B. L. Reactivation of Hippocampal Ensemble Memories During Sleep. Science 265, 676–679 (1994).

32. Fanselow, M. S. & Dong, H.-W. Are the Dorsal and Ventral Hippocampus Functionally Distinct Structures? Neuron 65, 7–19 (2010).

33. Strange, B. A., Witter, M. P., Lein, E. S. & Moser, E. I. Functional organization of the hippocampal longitudinal axis. Nat. Rev. Neurosci. 15, 655–669 (2014).

34. Hwang, H., Jin, S.-W. & Lee, I. Differential functions of the dorsal and intermediate regions of the hippocampus for optimal goal-directed navigation in VR space. eLife 13, RP97114 (2024).

35. Bast, T., Wilson, I. A., Witter, M. P. & Morris, R. G. M. From Rapid Place Learning to Behavioral Performance: A Key Role for the Intermediate Hippocampus. PLOS Biol. 7, e1000089 (2009).

36. Jarzebowski, P., Hay, Y. A., Grewe, B. F. & Paulsen, O. Different encoding of reward location in dorsal and intermediate hippocampus. Curr. Biol. 32, 834–841.e5 (2022).

37. Jin, S.-W. & Lee, I. Differential encoding of place value between the dorsal and intermediate hippocampus. Curr. Biol. 31, 3053–3072.e5 (2021).

38. Jin, S.-W., Ha, H.-S. & Lee, I. Selective reactivation of value- and place-dependent information during sharp-wave ripples in the intermediate and dorsal hippocampus. Sci. Adv. 10, eadn0416 (2024).

39. Okuyama, T., Kitamura, T., Roy, D. S., Itohara, S. & Tonegawa, S. Ventral CA1 neurons store social memory. Science 353, 1536–1541 (2016).

40. Oliva, A., Fernández-Ruiz, A., Leroy, F. & Siegelbaum, S. A. Hippocampal CA2 sharp-wave ripples reactivate and promote social memory. Nature 587, 264–269 (2020).

41. Watarai, A., Tao, K. & Okuyama, T. Representation of sex-specific social memory in ventral CA1 neurons. Science 389, eadp3814 (2025).

42. Boyle, L. M., Posani, L., Irfan, S., Siegelbaum, S. A. & Fusi, S. Tuned geometries of hippocampal representations meet the computational demands of social memory. Neuron 112, 1358–1371.e9 (2024).

43. Tsai, T.-C., Fang, Y.-S., Hung, Y.-C., Hung, L.-C. & Hsu, K.-S. A dorsal CA2 to ventral CA1 circuit contributes to oxytocinergic modulation of long-term social recognition memory. J. Biomed. Sci. 29, 50 (2022).

44. Kong, E., Lee, K.-H., Do, J., Kim, P. & Lee, D. Dynamic and stable hippocampal representations of social identity and reward expectation support associative social memory in male mice. Nat. Commun. 14, 2597 (2023).

45. Danjo, T., Toyoizumi, T. & Fujisawa, S. Spatial representations of self and other in the hippocampus. Science 359, 213–218 (2018).

46. Omer, D. B., Maimon, S. R., Las, L. & Ulanovsky, N. Social place-cells in the bat hippocampus. Science 359, 218–224 (2018).

47. Zhang, X. et al. Multiplexed representation of others in the hippocampal CA1 subfield of female mice. Nat. Commun. 15, 3702 (2024).

48. Dotson, N. M. & Yartsev, M. M. Nonlocal spatiotemporal representation in the hippocampus of freely flying bats. Science 373, 242–247 (2021).

49. Snyder, M. C., Qi, K. K. & Yartsev, M. M. Neural representation of human experimenters in the bat hippocampus. Nat. Neurosci. 27, 1675–1679 (2024).

50. Omer, D. B., Las, L. & Ulanovsky, N. Contextual and pure time coding for self and other in the hippocampus. Nat. Neurosci. 26, 285–294 (2023).

51. Ray, S. et al. Hippocampal coding of identity, sex, hierarchy, and affiliation in a social group of wild fruit bats. Science 387, eadk9385 (2025).

52. Forli, A. & Yartsev, M. M. Hippocampal representation during collective spatial behaviour in bats. Nature 621, 796–803 (2023).

53. O’Keefe, J. & Dostrovsky, J. The hippocampus as a spatial map. Preliminary evidence from unit activity in the freely-moving rat. Brain Res. 34, 171–175 (1971).

54. Moita, M. A. P., Rosis, S., Zhou, Y., LeDoux, J. E. & Blair, H. T. Hippocampal Place Cells Acquire Location-Specific Responses to the Conditioned Stimulus during Auditory Fear Conditioning. Neuron 37, 485–497 (2003).

55. Góis, Z. H. T. D. & Tort, A. B. L. Characterizing Speed Cells in the Rat Hippocampus. Cell Rep. 25, 1872–1884.e4 (2018).

56. Michon, F., Krul, E., Sun, J.-J. & Kloosterman, F. Single-trial dynamics of hippocampal spatial representations are modulated by reward value. Curr. Biol. 31, 4423–4435.e5 (2021).

57. Roux, L., Hu, B., Eichler, R., Stark, E. & Buzsáki, G. Sharp wave ripples during learning stabilize the hippocampal spatial map. Nat. Neurosci. 20, 845–853 (2017).

58. Michon, F., Sun, J.-J., Kim, C. Y., Ciliberti, D. & Kloosterman, F. Post-learning Hippocampal Replay Selectively Reinforces Spatial Memory for Highly Rewarded Locations. Curr. Biol. 29, 1436–1444.e5 (2019).

59. Huelin Gorriz, M., Takigawa, M. & Bendor, D. The role of experience in prioritizing hippocampal replay. Nat. Commun. 14, 8157 (2023).

60. Tingley, D. & Peyrache, A. On the methods for reactivation and replay analysis. Philos. Trans. R. Soc. B Biol. Sci. 375, 20190231 (2020).

61. Wu, W.-Y., Cheng, Y., Liang, K.-C., Lee, R. X. & Yen, C.-T. Affective mirror and anti-mirror neurons relate to prosocial help in rats. iScience 26, 105865 (2023).

62. Huang, Z. et al. Ventromedial prefrontal neurons represent self-states shaped by vicarious fear in male mice. Nat. Commun. 14, 3458 (2023).

63. Zhang, M., Wu, Y. E., Jiang, M. & Hong, W. Cortical regulation of helping behaviour towards others in pain. Nature 626, 136–144 (2024).

64. Sosa, M., Plitt, M. H. & Giocomo, L. M. A flexible hippocampal population code for experience relative to reward. Nat. Neurosci. 28, 1497–1509 (2025).

65. Gauthier, J. L. & Tank, D. W. A Dedicated Population for Reward Coding in the Hippocampus. Neuron 99, 179–193.e7 (2018).

66. Fenton, A. A. et al. Attention-Like Modulation of Hippocampus Place Cell Discharge. J. Neurosci. 30, 4613–4625 (2010).

67. Zeng, Y.-F. et al. Conjunctive encoding of exploratory intentions and spatial information in the hippocampus. Nat. Commun. 15, 3221 (2024).

68. Schuette, P. J. et al. Long-Term Characterization of Hippocampal Remapping during Contextual Fear Acquisition and Extinction. J. Neurosci. 40, 8329–8342 (2020).

69. Moita, M. A. P., Rosis, S., Zhou, Y., LeDoux, J. E. & Blair, H. T. Putting Fear in Its Place: Remapping of Hippocampal Place Cells during Fear Conditioning. J. Neurosci. 24, 7015–7023 (2004).

70. Dupret, D., O’Neill, J., Pleydell-Bouverie, B. & Csicsvari, J. The reorganization and reactivation of hippocampal maps predict spatial memory performance. Nat. Neurosci. 13, 995–1002 (2010).

71. Kentros, C. et al. Abolition of Long-Term Stability of New Hippocampal Place Cell Maps by NMDA Receptor Blockade. Science 280, 2121–2126 (1998).

72. Blair, G. J. et al. Hippocampal place cell remapping occurs with memory storage of aversive experiences. eLife 12, e80661 (2023).

73. Barnes, C. A., Suster, M. S., Shen, J. & McNaughton, B. L. Multistability of cognitive maps in the hippocampus of old rats. Nature 388, 272–275 (1997).

74. Agnihotri, N. T., Hawkins, R. D., Kandel, E. R. & Kentros, C. The long-term stability of new hippocampal place fields requires new protein synthesis. Proc. Natl. Acad. Sci. 101, 3656–3661 (2004).

75. Liberti, W. A., Schmid, T. A., Forli, A., Snyder, M. & Yartsev, M. M. A stable hippocampal code in freely flying bats. Nature 604, 98–103 (2022).

76. Wang, M. E., Yuan, R. K., Keinath, A. T., Ramos Alvarez, M. M. & Muzzio, I. A. Extinction of Learned Fear Induces Hippocampal Place Cell Remapping. J. Neurosci. 35, 9122–9136 (2015).

77. Monaco, J. D., Rao, G., Roth, E. D. & Knierim, J. J. Attentive scanning behavior drives one-trial potentiation of hippocampal place fields. Nat. Neurosci. 17, 725–731 (2014).

78. Takeuchi, T. et al. Locus coeruleus and dopaminergic consolidation of everyday memory. Nature 537, 357–362 (2016).

79. Kaufman, A. M., Geiller, T. & Losonczy, A. A Role for the Locus Coeruleus in Hippocampal CA1 Place Cell Reorganization during Spatial Reward Learning. Neuron 105, 1018–1026.e4 (2020).

80. Kim, J.-H., Choi, D.-E. & Shin, H.-S. The lateralized LC-NAergic system distinguishes vicarious versus direct fear in mice. Nat. Commun. 16, 2364 (2025).

81. McHugh, T. J. et al. Dentate gyrus NMDA receptors mediate rapid pattern separation in the hippocampal network. Science 317, 94–99 (2007).

82. Sahay, A. et al. Increasing adult hippocampal neurogenesis is sufficient to improve pattern separation. Nature 472, 466–470 (2011).

83. de Voogd, L. D. et al. The role of hippocampal spatial representations in contextualization and generalization of fear. NeuroImage 206, 116308 (2020).

84. Killcross, A. S., Kiernan, M. J., Dwyer, D. & Westbrook, R. F. Loss of latent inhibition of contextual conditioning following non-reinforced context exposure in rats. Q. J. Exp. Psychol. B 51, 75–90 (1998).

85. Bouton, M. E. Context and behavioral processes in extinction. Learn. Mem. 11, 485–494 (2004).

86. Pennartz, C. M. A. et al. The Ventral Striatum in Off-Line Processing: Ensemble Reactivation during Sleep and Modulation by Hippocampal Ripples. J. Neurosci. 24, 6446– 6456 (2004).

87. Baran, B., Pace-Schott, E. F., Ericson, C. & Spencer, R. M. C. Processing of Emotional Reactivity and Emotional Memory over Sleep. J. Neurosci. 32, 1035–1042 (2012).

88. Braun, E. K., Wimmer, G. E. & Shohamy, D. Retroactive and graded prioritization of memory by reward. Nat. Commun. 9, 4886 (2018).

89. Singer, A. C. & Frank, L. M. Rewarded Outcomes Enhance Reactivation of Experience in the Hippocampus. Neuron 64, 910–921 (2009).

90. Ambrose, R. E., Pfeiffer, B. E. & Foster, D. J. Reverse Replay of Hippocampal Place Cells Is Uniquely Modulated by Changing Reward. Neuron 91, 1124–1136 (2016).

91. Patel, J., Schomburg, E. W., Berényi, A., Fujisawa, S. & Buzsáki, G. Local Generation and Propagation of Ripples along the Septotemporal Axis of the Hippocampus. J. Neurosci. 33, 17029–17041 (2013).

92. Morici, J. F., Silva, A., Lima-Paiva, I., Pronier, É. & Girardeau, G. Dorsoventral hippocampus neural assemblies reactivate during sleep following an aversive experience. Nat. Neurosci. 29, 1133–1144 (2026).

93. Girardeau, G., Cei, A. & Zugaro, M. Learning-Induced Plasticity Regulates Hippocampal Sharp Wave-Ripple Drive. J. Neurosci. 34, 5176–5183 (2014).

94. Han, Y. et al. Bidirectional cingulate-dependent danger information transfer across rats. PLoS Biol. 17, e3000524 (2019).

95. Terranova, J. I. et al. Hippocampal-amygdala memory circuits govern experience-dependent observational fear. Neuron S0896-6273(22)00058–7 (2022) doi:10.1016/j.neuron.2022.01.019.

96. Han, Y., Sichterman, B., Maria, C., Gazzola, V. & Keysers, C. Similar levels of emotional contagion in male and female rats. Sci. Rep. 10, 2763 (2020).

97. Kim, A., Keum, S. & Shin, H.-S. Observational fear behavior in rodents as a model for empathy. Genes Brain Behav. 18, e12521 (2019).

98. van Daal, R. J. J. et al. Implantation of Neuropixels probes for chronic recording of neuronal activity in freely behaving mice and rats. Nat. Protoc. 16, 3322–3347 (2021).

99. Almonte, A. et al. Enhanced ventral hippocampal synaptic transmission and impaired synaptic plasticity in a rodent model of alcohol addiction vulnerability. Sci. Rep. 7, (2017).

100. Pereira, T. D. et al. SLEAP: A deep learning system for multi-animal pose tracking. Nat. Methods 19, 486–495 (2022).

101. Schlingloff, D., Káli, S., Freund, T. F., Hájos, N. & Gulyás, A. I. Mechanisms of sharp wave initiation and ripple generation. J. Neurosci. Off. J. Soc. Neurosci. 34, 11385– 11398 (2014).

102. Buzsáki, G. Hippocampal sharp wave-ripple: A cognitive biomarker for episodic memory and planning. Hippocampus 25, 1073–1188 (2015).

103. Bürkner, P.-C. brms: An *R* Package for Bayesian Multilevel Models Using *Stan*. J. Stat. Softw. 80, (2017).

104. Makowski, D., Ben-Shachar, M. S. & Lüdecke, D. bayestestR: Describing Effects and their Uncertainty, Existence and Significance within the Bayesian Framework. J. Open Source Softw. 4, 1541 (2019).

105. Kass, R. E. & Raftery, A. E. Bayes Factors. J. Am. Stat. Assoc. 90, 773–795 (1995).

106. Keysers, C., Gazzola, V. & Wagenmakers, E.-J. Using Bayes factor hypothesis testing in neuroscience to establish evidence of absence. Nat. Neurosci. 23, 788–799 (2020).

