## Supplementary material for "The Intermediate Hippocampus Integrates Shock-Observation and Spatial Information during Observational Fear Memory in Male Rats"

| | Shock Obs | count | | | percentage | | |
|---|---|---|---|---|---|---|---|
| subregion | area | excited | inhibited | unresponsive | excited | inhibited | unresponsive |
| dorsal | CA1 | 4 | 2 | 26 | 12.5 | 6.25 | 81.25 |
| | CA2 | 0 | 1 | 2 | 0 | 33.33 | 66.67 |
| | CA3 | 1 | 2 | 35 | 2.63 | 5.26 | 92.11 |
| intermediate | CA1 | 10 | 3 | 54 | 14.93 | 4.8 | 80.60 |
| | CA3 | 13 | 6 | 58 | 16.88 | 7.79 | 75.33 |
| ventral | CA1 | 2 | 6 | 40 | 4.17 | 12.5 | 83.33 |
| | CA3 | 4 | 8 | 60 | 5.56 | 11.11 | 83.33 |

Supplementary Table 1: Quantification of pyramidal neurons by estimated recording location and response at the moment of shock observation.

| | Shock Obs | count | | | percentage | | |
|---|---|---|---|---|---|---|---|
| subregion | area | excited | inhibited | unresponsive | excited | inhibited | unresponsive |
| dorsal | CA1 | 3 | 0 | 5 | 37.5 | 0 | 62.5 |
| | CA2 | 1 | 0 | 0 | 100 | 0 | 0 |
| | CA3 | 4 | 2 | 10 | 25 | 12.5 | 62.5 |
| intermediate | CA1 | 6 | 0 | 4 | 60 | 0 | 40 |
| | CA3 | 7 | 5 | 17 | 24.14 | 17.24 | 58.62 |
| ventral | CA1 | 15 | 8 | 32 | 27.27 | 14.55 | 58.18 |
| | CA3 | 11 | 4 | 15 | 36.67 | 13.33 | 50 |

Supplementary *Table 2: Quantification of interneurons by estimated recording location and response at the moment of shock observation.*

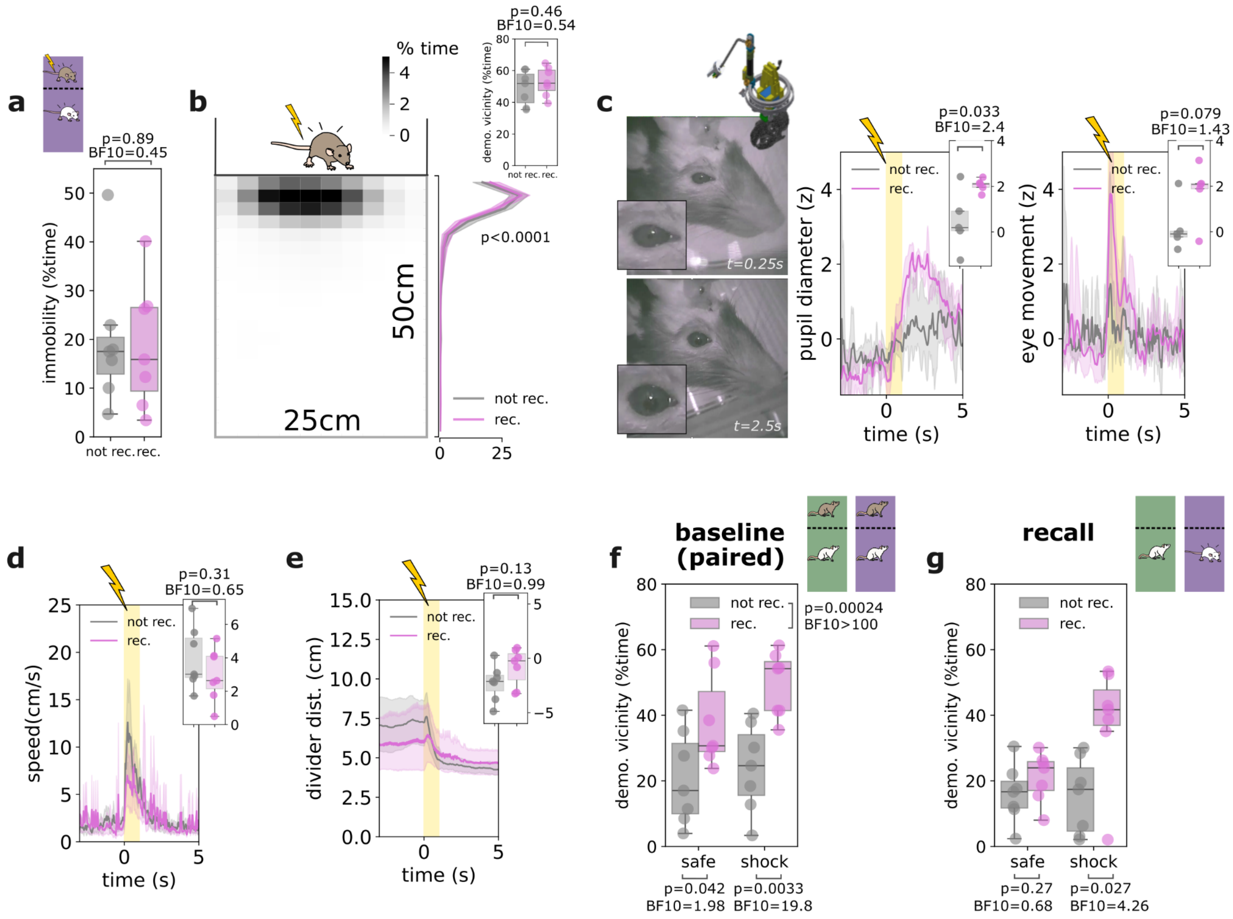


***Supplementary Figure1: Behavioral correlates of contextual vicarious fear learning***

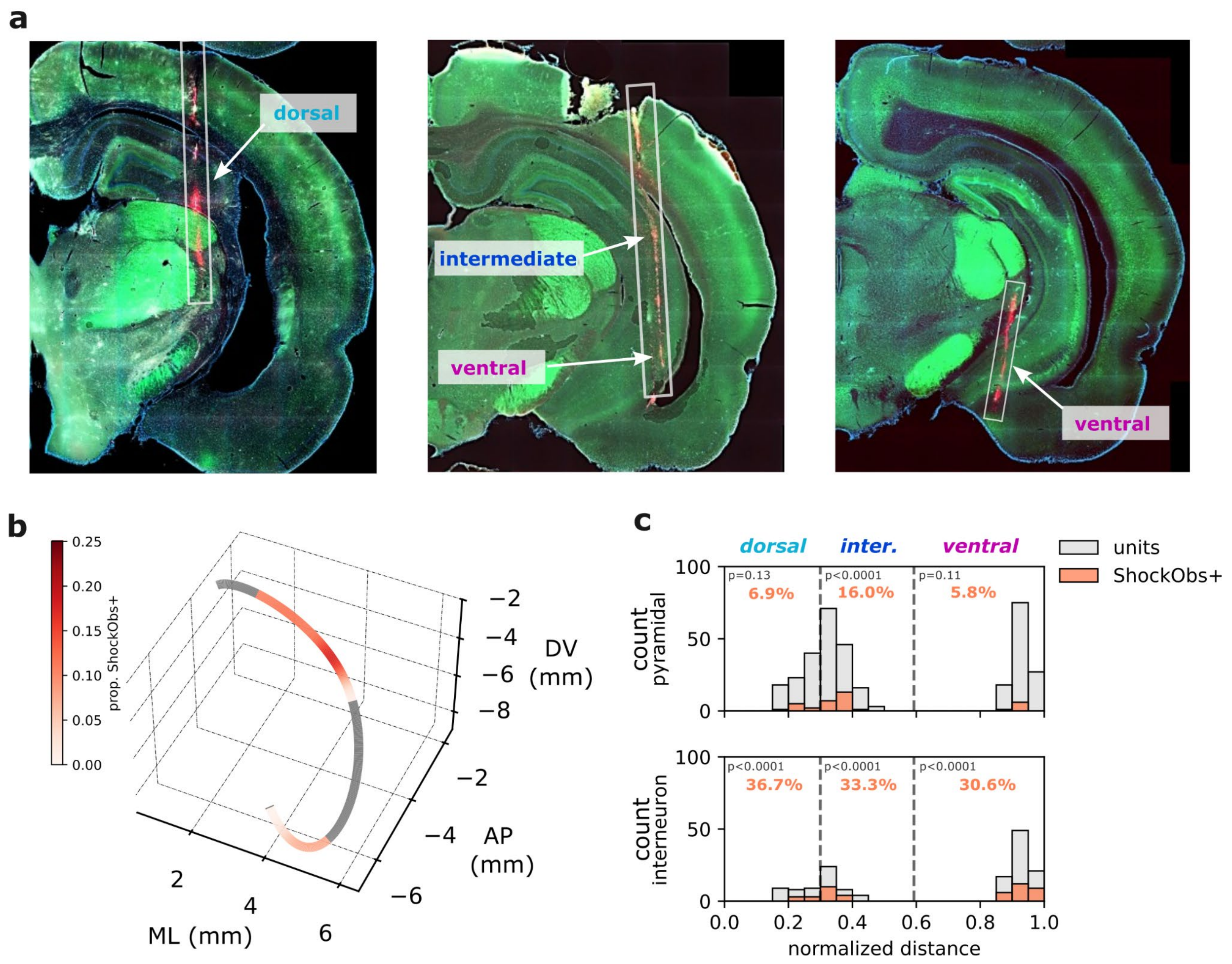


**Supplementary Figure2: Anatomical mapping of hippocampal recording sites**

**a.** Example coronal sections used to reconstruct implantation sites, with probe shank tracks crossing the hippocampal principal sheets at coordinates classified as corresponding to the dorsal, intermediate or ventral hippocampus. Images were processed in QuPath (v0.2.3), where each color channel was thresholded individually. **b.** Proportion of shock observation responsive pyramidal neurons mapped onto a 3-D reconstruction of the hippocampal septo-temporal axis along mid-area CA1. Gray segments indicate portions of the septotemporal axis that were not sampled. **c.** Distribution of all (units, in grey) pyramidal (top panel, N = 337 neurons) or interneurons (bottom panel, N = 149) and shock observation responsive units ( shockobs+, orange) along the septotemporal axis as shown in B. Dashed lines mark the border used to define dorsal, intermediate and ventral hippocampal subregions.

P values are two-sided Monte Carlo P values for testing whether the number of responsive neurons exceeds that expected by chance estimated from random temporal shifting. Source data are provided as a Source Data file.

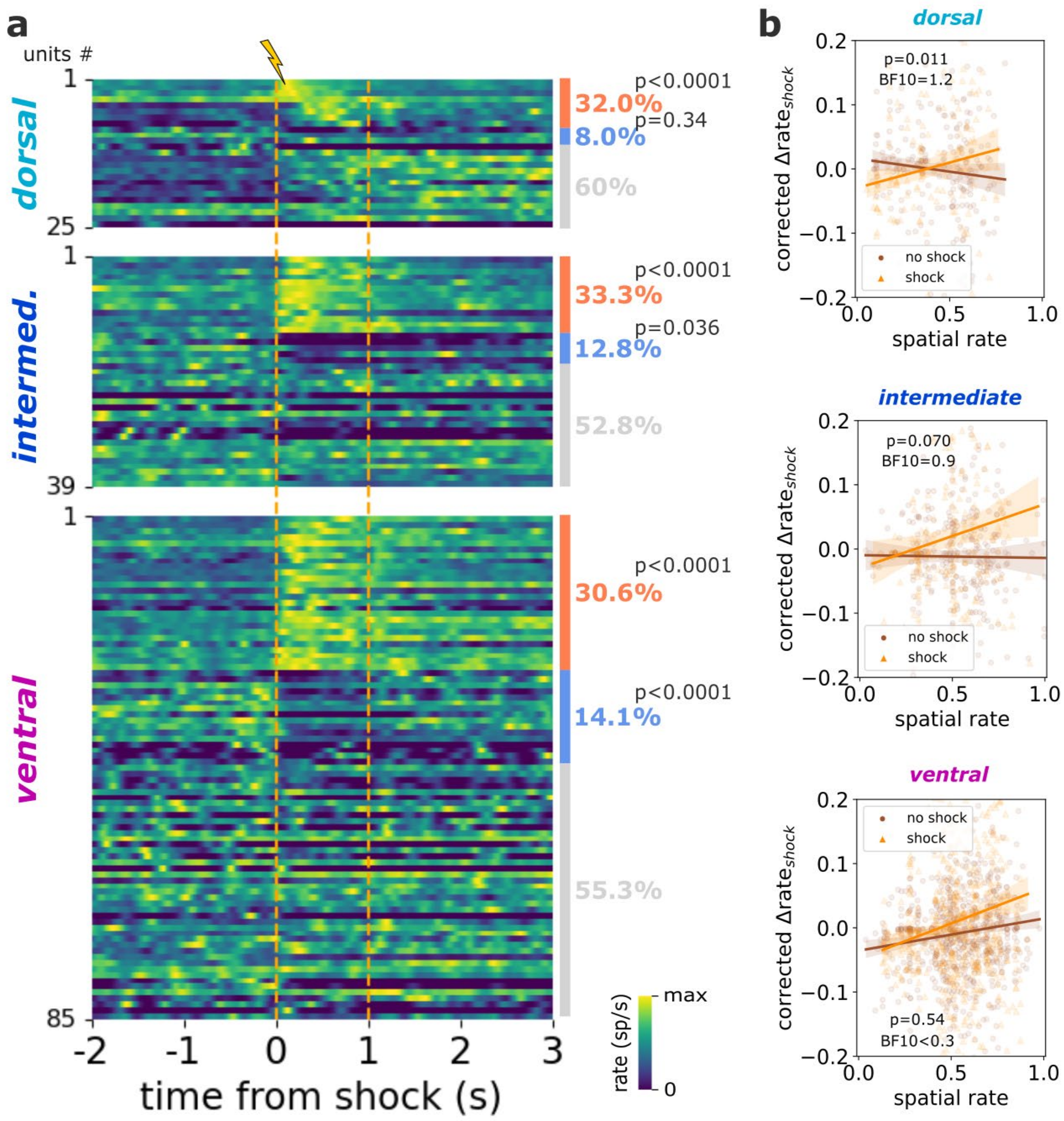


**Supplementary Figure3: Interneurons response to shock observation**

**a.** Heatmap representing the mean firing rate of single putative interneuron split by hippocampal sub-region (dorsal, intermediate, ventral). For each sub-region, neurons are ordered from top to bottom as follows: significantly excited (orange), significantly inhibited (blue) and finally not significantly modulated (grey) single neurons. Overall percentages are indicated on the right side of the heatmaps. Orange dashed vertical lines represent the onset and offset of footshock delivery. **b.** For each hippocampal subregion, linear regression between corrected neuronal response for speed and distance to the demonstrator's compartment and estimated basal spatial firing rate at times of footshock delivery (gold) or matched control-moments without shocks (sienna).

In (**a**) Monte Carlo P value for testing whether the number of responsive neurons exceeds that expected by chance estimated from random temporal shifting. In (**b**), mean regression line with 95% confidence interval. Dots represent single trial responses. P value and $BF_{10}$ for the linear mixed model's fixed effect of the interaction: baseline spatial rate X footshock delivery times. Source data are provided as a Source Data file.

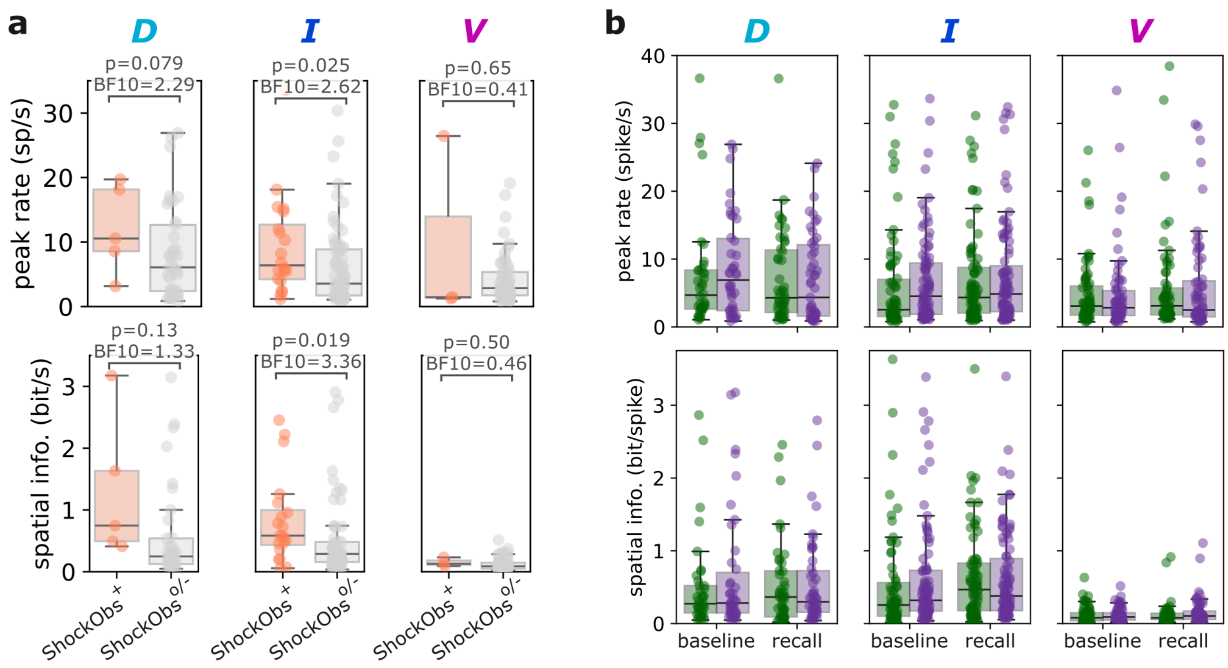


**Supplementary Figure4: Spatial firing properties**

**a.** Spatial tuning properties of putative pyramidal neurons during solo baseline split by hippocampal subregion (dorsal, D, intermediate, I, and ventral, V) separately for neurons significantly excited ($ShockObs^{+}$, in orange, N = 5/23/6 neurons in D/I/V) or not (unresponsive and inhibited, $ShockObs^{o/-}$, in grey, N = 68/121/114 neurons in D/I/V) by the shock observation. Top row, maximum field rate. Bottom row, spatial information. **b.** For all putative pyramidal neurons, peak field rate (top) and spatial information (bottom) for spatial rate maps estimated from the exploration of the observer's compartment in the safe (green) and shock (purple) context, and during solo baseline and recall.

Boxplots center line represents the median, the box limits the 25th and 75th percentiles, and the whiskers extend to values up to 1.5 x the interquartile range; points indicate values for individual neurons. In **a**, uncorrected P values and $BF_{10}$ from two-sided student T-test. In **b,** Top, two-way ANOVA (context × phase), in dorsal: context ($F(1,258)=0.74$, $p=0.39$, $\eta^2p=0.003$), phase ($F(1,258)=0.17$, $p=0.68$, $\eta^2p<0.001$), interaction ($F(1,258)=0.02$, $p=0.88$, $\eta^2p<0.001$); in intermediate: context ($F(1,543)=3.67$, $p=0.056$, $\eta^2p=0.007$), phase ($F(1,543)=2.96$, $p=0.086$, $\eta^2p=0.005$), interaction ($F(1,543)=0.02$, $p=0.88$, $\eta^2p<0.001$); context ($F(1,417)=0.08$, $p=0.77$, $\eta^2p<0.001$), phase ($F(1,417)=0.19$, $p=0.67$, $\eta^2p<0.001$), or interaction ($F(1,417)=0.89$, $p=0.35$, $\eta^2p=0.002$). Bottom, two-way ANOVA (context × phase), in dorsal: context ($F(1,276)=0.67$, $p=0.41$, $\eta^2p=0.002$), phase ($F(1,276)=0.88$, $p=0.35$, $\eta^2p=0.003$), interaction ($F(1,276)=0.29$, $p=0.59$, $\eta^2p=0.001$); in intermediate: context ($F(1,572)=0.31$, $p=0.58$, $\eta^2p<0.001$), phase ($F(1,572)=2.44$, $p=0.12$, $\eta^2p=0.004$), interaction ($F(1,572)=0.32$, $p=0.57$, $\eta^2p<0.001$); in ventral: context ($F(1,476)=1.05$, $p=0.31$, $\eta^2p=0.002$), phase ($F(1,476)=0.58$, $p=0.45$, $\eta^2p=0.001$), or interaction ($F(1,476)=0.81$, $p=0.37$, $\eta^2p=0.002$). Source data are provided as a Source Data file.

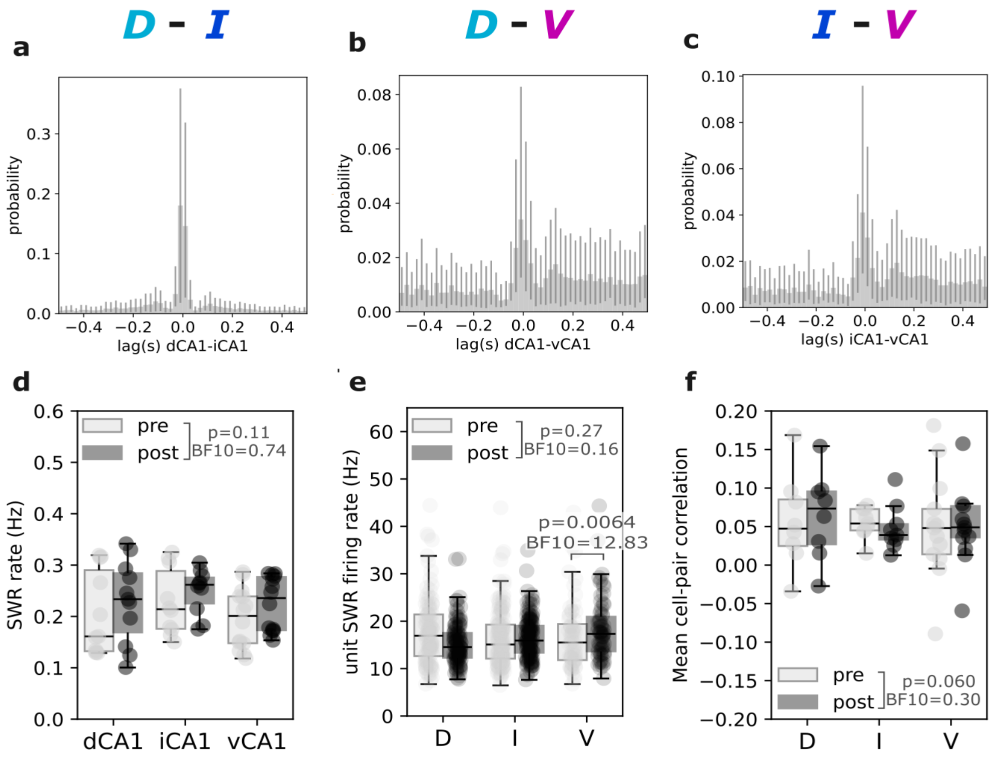


**Supplementary Figure5: SWR-related activity across hippocampal subregions**

Stronger dorsal-intermediate SWR synchrony compared to dorsal-ventral and intermediate-ventral SWR occurrences during the post-rest phase. Mean cross-correlograms for the occurrence of SWRs detected in the dorsal and intermediate hippocampus (N = 7 rats) (**a**), dorsal and ventral hippocampus (N = 11 rats) (**b**), and intermediate and ventral hippocampus (N = 10 rats) (**c**). **d.** Mean rates of sharp-wave ripples occurrence detected in the dorsal (dCA1, N = 11 rats), intermediate (iCA1, N=10) and ventral (vCA1, N = 14 rats) hippocampus, computed for each rat during the pre-shock (light grey) and post-shock (dark grey) rest phases. **e.** Mean single-unit firing rate of recorded pyramidal neurons during sharp-wave ripple occurrences detected in the pre-shock and post-shock rest phases, and separated for units recorded from the dorsal (D, N= 73 neurons), intermediate (I, N = 144 neurons) and ventral (V, N = 120 neurons) hippocampus. **f.** For each rat, the mean pyramidal cell pair firing rate correlation during sharp-wave ripples detected in the pre-shock and post-shock rest phases, and separated for units recorded from the dorsal (D, N = 8 rats), intermediate (I, N = 9 rats) and ventral (V, N = 13 rats) hippocampus.

Error bars represent the 95% confidence interval estimated from 500 bootstrapping iterations. In **d-f**, Boxplots center line represents the median, the box limits the 25th and 75th percentiles, and the whiskers extend to values up to 1.5 x the interquartile range; points indicate values for individual animals. In **d**, two-way ANOVA (subregion × sleep phase): subregion ($F(2,64)=1.13$, $p=0.33$, $\eta^2p=0.034$), phase ($F(1,64)=2.51$, $p=0.12$, $\eta^2p=0.038$), interaction ($F(2,64)=0.02$, $p=0.98$, $\eta^2p<0.001$). In **e**, two-way ANOVA (subregion × sleep phase): subregion ($F(2,660)=2.27$, $p=0.10$, $\eta^2p=0.007$), phase ($F(1,660)=1.25$, $p=0.26$, $\eta^2p=0.002$), interaction ($F(2,660)=5.12$, $p=0.006$, $\eta^2p=0.015$). In **f**, two-way ANOVA (subregion × sleep phase): phase ($F(1,54)=0.27$, $p=0.61$, $\eta^2p=0.005$), subregion ($F(2,54)=0.11$, $p=0.89$, $\eta^2p=0.004$), interaction ($F(2,54)=0.23$, $p=0.80$, $\eta^2p=0.008$). In **d-f**, corrected P value and Bayesian $BF_{10}$ from post hoc ANOVA student T-test are reported. Source data are provided as a Source Data file.
